# Building a robust backbone for *Astragalus* (Fabaceae) using a clade-specific target enrichment bait set

**DOI:** 10.1101/2024.11.19.624276

**Authors:** Daniele Buono, Gudrun Kadereit, Aaron Liston, Shahin Zarre, Diego F. Morales-Briones

## Abstract

**Premise of the study:** With over 3100 species, *Astragalus* (Fabaceae) has long fascinated botanists as the largest genus of flowering plants. With an origin in the Middle Miocene, *Astragalus* has one of the highest diversification rates known in flowering plants. Comprehensive taxonomic treatments exist, and the genus is currently subdivided into 136 sections in the Old World and 93 sections in the New World based on morphological characters. Despite considerable efforts, a comprehensive and well-resolved phylogeny of the genus is still lacking.

**Methods:** Here, we reconstruct the backbone phylogeny of *Astragalus* using a custom bait set capturing 819 loci specifically designed for a target enrichment approach in the Astragalean clade. We carefully selected a set of 107 taxa representing all major clades currently recognized in *Astragalus*. Of those, 80 newly sequenced taxa were obtained from herbarium specimens as old as 110 years.

**Key results:** We retrieved all the targeted loci and additional off-target plastome sequences for all samples, including the 80 herbarium specimens. Our phylogenetic analysis reinforced the currently accepted backbone phylogeny of *Astragalus* with high support and unprecedented details, additionally providing insights into cytonuclear phylogenetic conflicts in the genus. Evidence for potential reticulate evolution was found, providing a possible explanation for the conflicts observed.

**Conclusions:** This work represents an important milestone in obtaining a comprehensive, herbarium-based phylogeny of *Astragalus*, which will constitute the base to study a wealth of relevant biological questions, for example, the still unanswered question of what drove the rapid diversification of *Astragalus*, with important repercussions on our understanding of diversification in natural contexts.

## Introduction

*Astragalus* L. (Fabaceae, subfamily Papilionoideae), with about 3,100 species (POWO, 2024), is considered the largest genus of flowering plants and represents a striking example of rapid radiation (Azani et al., 2019; Moonlight et al., 2024). The genus has a nearly worldwide distribution, occurring mainly in semi-arid regions of the Northern Hemisphere, with about 2,600 species distributed in the Old World (Eastern Hemisphere) and about 500 species in the New World (Western Hemisphere; Maassoumi and Ashouri, 2022). The size of the distribution areas of *Astragalus* species ranges from continentally widespread (13 species have a circumboreal distribution) to hundreds of narrow endemics (Azani et al., 2019). The center of diversity of the genus is cool to warm arid or semiarid and mountainous regions, especially in southwest Asia in the Irano-Turanian floristic region, which hosts more than 1,500 species (Soltani et al., 2021). Iran displays the highest rate of endemism, with about 620 endemic species out of 900 in the country (Maassoumi and Ashouri, 2022). *Astragalus* includes morphologically diverse groups of annuals (∼120 spp.) to perennial herbs (∼2,500 spp.) and spiny cushions (∼300 spp.). Despite its ubiquity and striking diversity, the genus is relatively poorly studied, and its phylogeny is yet to be explored comprehensively (Hardion et al., 2016; but see Azani et al., 2017, 2019; Folk et al., 2024, discussed below).

### Species diversity in *Astragalus*

*Astragalus* does not show higher rates of diversification than its relatives in the Astragalean clade (e.g., genus *Oxytropis* DC. with about 600 species; POWO, 2024) or compared to other temperate papilionoid legumes. Instead, the Astragalean clade (sensu Sanderson and Liston, 1995) as a whole (ca. 3,900 species) is significantly more diverse than its closest relatives (e.g., *Caragana* Lam., *Colophaca* Fisch., *Chesneya* Lindl. ex Engl., and *Guedenstaedtia* Fisch.) (Sanderson and Wojciechowski, 1996; Koenen et al., 2013). There seems to be no distinctive ecological trait in the Astragalean clade that could be regarded as or correlated with any obvious key innovation that would explain the diversification observed. Therefore, Sanderson and Wojciechowski (1996) hypothesized that demographic factors, such as population fragmentation and isolation, may better explain diversification within this group. Furthermore, ecological specialization in *Astragalus* to edaphic conditions and extreme microhabitats unfavorable for most other plants suggests that adaptation to local conditions may lead to divergence and persistence of lineages (Rundel et al., 2015; Kenicer, 2005; Maassoumi and Khajoei Nasab, 2023). Scherson et al. (2008) indicated that new ecological opportunities caused by the formation of different environments and microclimates as a result of the uplift of the Andes during the Pliocene (2–4 Ma) promoted the high diversification rates of *Astragalus* observed in the Southern Andes. Interestingly, the species diversity of *Astragalus* decreases towards the Northern Andes, with no records of the genus found north of Ecuador. Similarly, Azani et al. (2019) hypothesize that the intense uplift of the Qinghai-Tibetan Plateau (QTP) during the Late Miocene resulted in prolonged droughts promoting the formation of semi-desert habitats, which might have favored the diversification of *Astragalus*. On the other hand, Hardion et al. (2016) supported the hypothesis that the main driver of diversification in the Mediterranean xerophytic *Astragalus* sect. *Tragacantha* DC. is range fragmentation rather than a coastal-mountain ecological shift. The underlying causes of the high diversity might be a mix of common intrinsic traits and different environmental and geographic settings among clades and regions, which makes a macroevolutionary study of this fascinating genus highly desirable.

The origin of the genus has been dated to the Middle Miocene 16 Mya (12.27–20.76 Mya) and placed in West Asia based on nrDNA (ITS) and cpDNA (*trnK/matK* + *ycf1*) data (Azani et al., 2019). A rapid diversification followed, starting from about 14 Mya, with subsequent range expansion repeatedly in the Mediterranean area and then North America (Azani et al., 2019). There exists a strong relationship between chromosome number and geographic distribution. Most Old World and circumboreal species have euploid chromosome numbers based on *n* = 8, with *n* = 8, 16, 32, 48, 64, 96, with common polyploids. Almost all American species, except approximately 15 North American species, form the Neo-*Astragalus* clade characterized by an aneuploid chromosome number of *n* = 11, 12, 13, 14, 15 (Wojciechowski et al., 1993, 1999). The origin of the aneuploid American *Astragalus* is still unresolved, with Azani et al. (2017) suggesting a Mediterranean origin of around 4.36 Mya, while the study of Su et al. (2021) supported an Asian origin instead. Azani et al. (2019) suggested multiple origins of New World *Astragalus* from the Diholcos clades for the aneuploid species and from the Hamosa, Phaca, and Hypoglottis clades for the euploid New World species. In contrast, Folk et al. (2024) suggested that a single dispersal event from western Asia about 9.8 Mya, thus significantly earlier than previous estimates, is behind the broad ancestral distribution across the Americas.

### Current subgeneric classifications and their limitations

*Astragalus* species are classified into 136 sections in the Old World (Podlech and Zarre, 2013) and 93 sections in the New World (Barneby, 1964) based on morphological characters. Molecular phylogenetic studies demonstrated that the most important morphological features used in classical taxonomic studies (e.g., medifixed vs. basifixed indumentum, perennial vs. annual) are homoplasious and have originated several times independently (Zarre and Azani, 2012). In *Astragalus*, 11 major clades were recovered in a phylogenetic tree by Azani et al. (2017) based on nuclear ITS + plastid *trnK*/*matK*. Su et al. (2021), based on 65 CDS of 117 plastomes, recovered an additional clade and referred to it as Pseudosesbanella. Otherwise, their phylogenetic tree mostly overlapped with that of Azani et al. (2017), even though it lacked samples from the Ophiocarpus clade. Folk et al. (2024) published the most recent molecular phylogenetic study, including a larger New World *Astragalus* taxon sampling. The authors included about 900 species (373 taxa belonging to the New World, corresponding to ∼45% of the total, and 474 to the Old World, corresponding to ∼20% of the total). Their phylogeny was based on a selection of approximately 100 loci intended to cover the entire nitrogen-fixing clade. The authors recovered a phylogeny with most backbone nodes having high support except for the Astragalean clade (local posterior probability [LPP] = 0.08) and very low support and poorly resolved relationships within the main clades (average LPP = 0.605). Several inconsistencies with currently accepted phylogenies of the genus were observed. The Glottis clade (sensu Azani et al., 2017), for which only one species was sampled (*A. epiglottis* = *Biserulla epiglottis*), was placed outside Eu-Astragalus. The position of Neo-*Astragalus* conflicts with all previous molecular studies, placing it as an independent clade along the backbone instead of nested within the Diholcos clade. Furthermore, the Contortuplicata clade (sensu Azani et al., 2017) was ambiguously placed in a polytomy with the Hamosa clade (sensu Azani et al., 2017). Additionally, even though the clades recovered in their phylogeny largely aligned with Azani et al. (2017) and Su et al. (2021), the authors adhered to an outdated and less informative clade naming system (Kazempour Osaloo et al., 2003, 2005).

Those conflicts were considered to be the result of different evolutionary properties of plastid loci and nrDNA compared with low-copy nuclear genes, advocating for further studies to investigate incongruences between nuclear and organellar signals (Folk et al., 2024). Therefore, relationships among major clades inside *Astragalus* and the tempo of their evolution remain unclear, highlighting the need for further molecular studies based on additional data.

### Target enrichment

The use of low-copy or single-copy nuclear genes (SCN) is highly beneficial for plant phylogenetics because they offer more rapidly evolving characters compared to regularly applied chloroplast and ribosomal sequences (Zimmer and Wen, 2015). To obtain those sequences, short-read sequencing techniques, such as target enrichment, are useful when working with dried plant material from herbaria, which usually contain short DNA fragments (often less than 100bp; Forrest et al., 2019). Target enrichment can be used with highly fragmented gDNA to obtain high coverage of targeted loci (Andermann et al., 2020; McKain et al., 2018). This method uses short RNA probes (usually 80–120 bp) that hybridize into complementary sequence library fragments (McKain et al., 2018). Hybridized fragments can be bound to magnetic beads and separated from the rest of the library, improving the depth of coverage for genes of interest. This characteristic makes the method useful for analyzing specific variants, sequence exomes, and large numbers of genes, inferring genome duplications and ortholog genes. Probe design was a limitation of the target enrichment technique in the past and was originally focused on highly conserved loci to obtain broad taxonomic coverage (Faircloth et al., 2012; Pezzini et al., 2023). However, thanks to the increasing availability of transcriptome sequence data across all major clades, it has become more accessible to design custom probe sets for a specific plant group (McKain et al., 2018; Vatanparast et al., 2018). Taxon-specific probe sets usually work better than universal ones because they are less biased towards more conserved regions and provide higher resolution at a finer taxonomic scale, recovering more genomic variation (Pezzini et al., 2023).

### Goal of the study

In this study, we attempt to resolve subgenus-level relationships within *Astragalus* by using a bait set for target enrichment, which includes 819 loci derived from transcriptome data of Astragalean taxa. The bait set was tested with DNA samples extracted from herbarium specimens as old as 110 years, covering many of the sections currently supported by molecular data in the genus, plus several species representing other genera in the Astragalean clade. This newly produced data set was integrated with publicly available published transcriptome data. A backbone phylogeny was reconstructed and compared with currently accepted and newly proposed phylogenies of *Astragalus* (Azani et al., 2017; Azani et al., 2019; Folk et al., 2024). The possibility of reconstructing an organelle phylogeny using off-target regions was also explored, providing unprecedented insights into cytonuclear phylogenetic conflicts in the Astragalean clade.

## MATERIAL AND METHODS

### Bait-set design

We designed a custom bait set for the Astragalean clade (*sensu* Zhao et al., 2021). We identified putative SCN genes with MarkerMiner v.1.2 (Chamala et al., 2015) with default settings, except the minimum transcript length set to 500 bp. We used the transcriptomes of 17 species (eight from *Astragalus*) from 10 genera across the Astragalean clade (Zhao et al., 2021; Table 1) and the genomes of *Cicer arietinum* L. (Varshney et al., 2013), *Medicago truncatula* Gaertn. (Tang et al., 2014), and *Trifolium pratense* L. (De Vega et al., 2015) as references. SCN genes identified with MarkerMiner were further filtered and split into exons using GoldFinder (Vargas et al., 2019), requiring loci with at least 500 bp and coverage of at least three species. Individual exons fasta files were realigned with MACSE v.2.06 (Ramwez et al., 2018) to correct codon frames and visually inspected in GENEIOUS v.11.1.5 (https://www.geneious.com) to remove potential misassemblies from the beginning or ends of the alignments. Additionally, exon alignments were filtered to have at least 500 bp and pairwise identity between 75% and 98%. This resulted in 819 exons from 686 genes across the Astragalean clade. For bait design, we filtered the 819 exon alignments to include only two sequences, one from a species of *Astragalus* and one from a species from any other genus in the Astragalean clade. The custom set of 80 bp biotinylated RNA baits (myBaits) was manufactured by Daicel Arbor Biosciences (Ann Arbor, MI, USA) with a 2× tiling density.

**Table 1.**
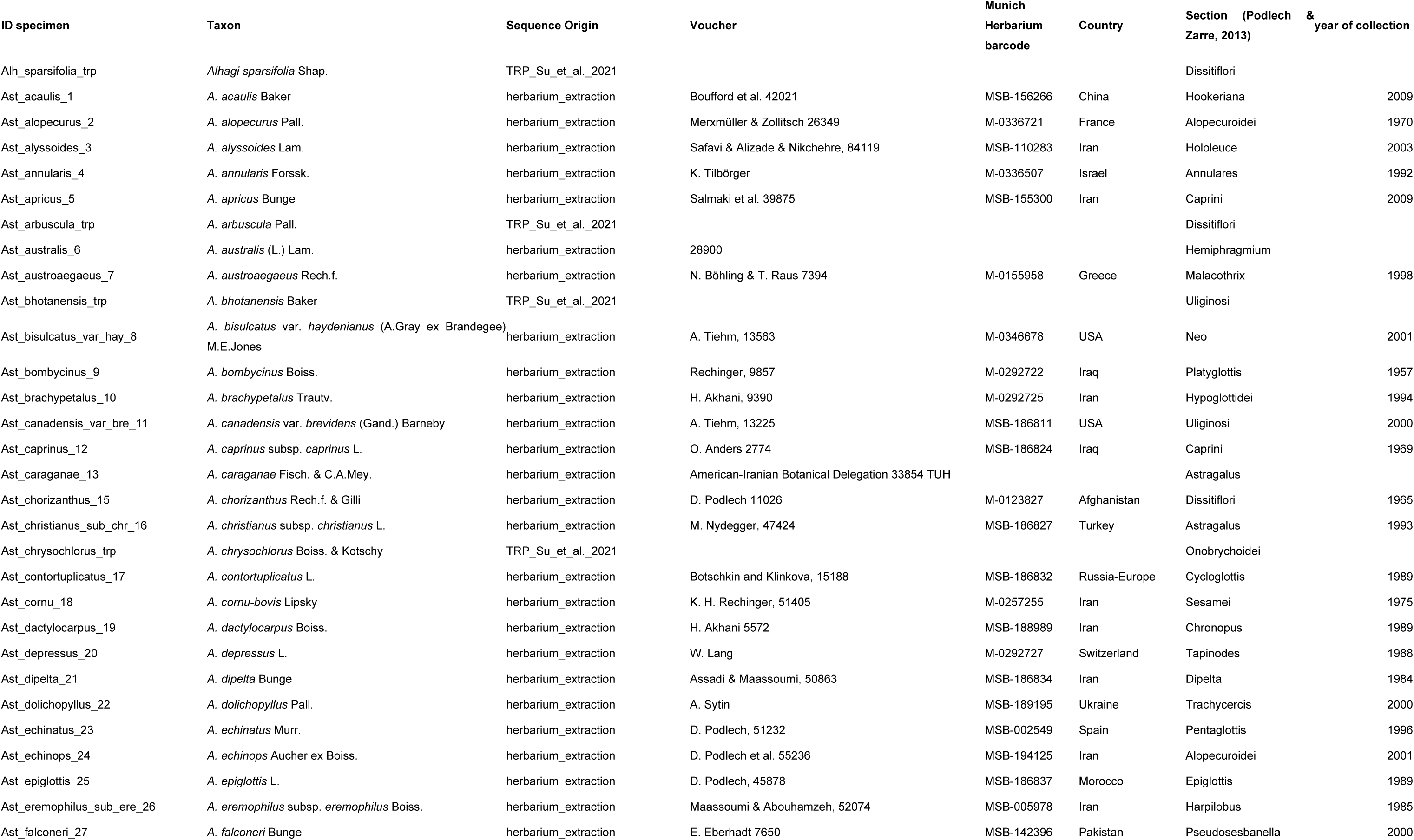
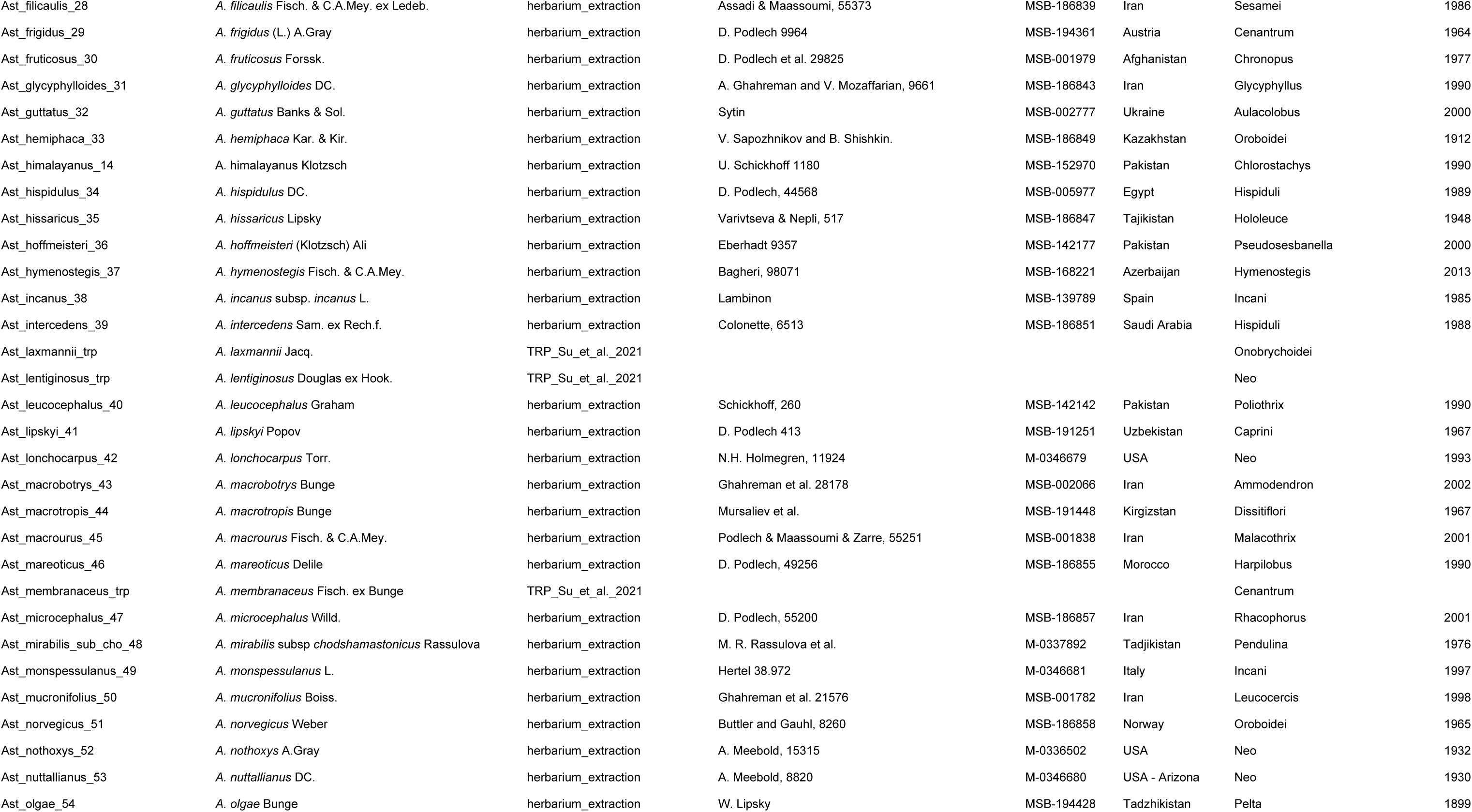
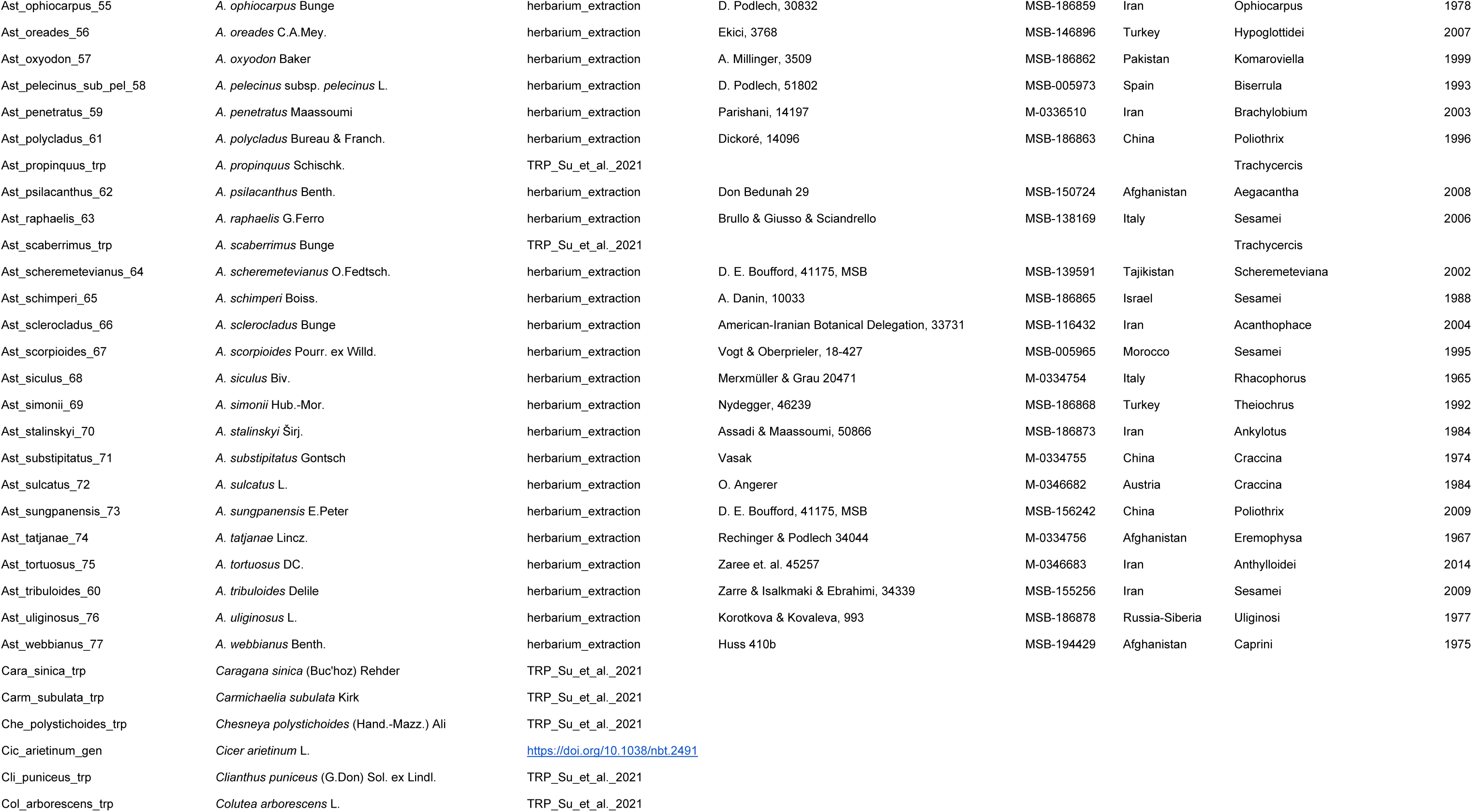
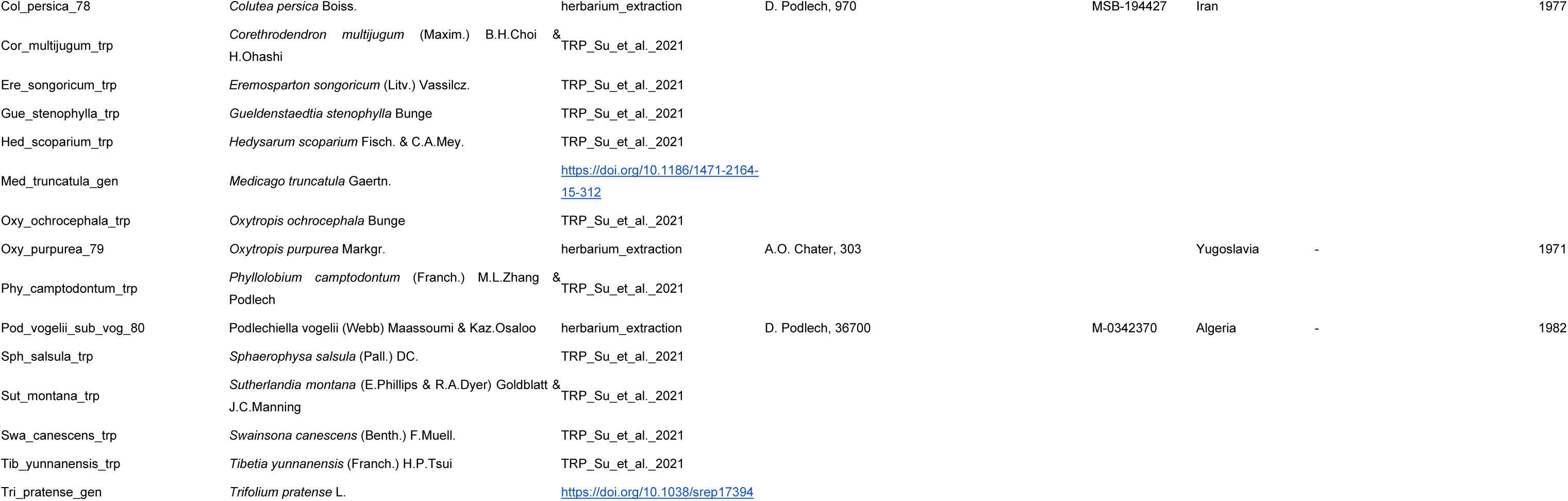
Samples used in this study. In the column Sequence Origin, hebarium_extraction indicates newly generated sequences, TRP_Su_et_al._2021 transcriptome data obtained from Su et al. (2021), and specific study when DOI was provided.

### Taxon sampling

To compare our phylogeny with previous ones and to cover the major clades recognized in *Astragalus*, we chose to sample species from the major clades obtained in the phylogenies of Azani et al. (2017), Su et al. (2021), and Folk et al. (2024). When specimens were available at the Botanical State Collection Munich (herbaria M and MSB), sampled vouchers were identical to those of Azani et al. (2017, Table S1). To increase the representativeness of the Astragalean clade and add outgroups, additional taxa, of which transcriptome (Zhao et al., 2021) and genomic data were available online, were added to our dataset (Table 1).

### Library preparation, target enrichment, and sequencing

DNA was extracted from ∼20 mg of dry herbarium plant material using the NucleoSpinⓇ Plant II kit (Macherey-Nagel, Düren, Germany), following the manufacturer’s manual with slight modifications. Cell lysis in step 2a was performed using 600 μl buffer PL1, no RNase A was added since the plant material is too old to preserve RNA, and incubation lasted for 1:30h. An extra washing was performed in step 6, adding 350 μl buffer PW2 before drying the membrane completely. Finally, DNA was eluted in a 50 μl buffer PE (5 mM Tris/HCl, pH 8.5). DNA concentration was measured using an Invitrogen™ Qubit™ 4 Fluorometer using the High Sensitivity (HS) assay kit (Thermo Fisher Scientific, Waltham, Massachusetts, USA), and fragmentation was visually evaluated on 1% agarose gel.

Before library preparation, genomic DNA was diluted to 250 ng in 55 μl buffer PE and sonicated using a Covaris M220 Focused-ultrasonicator (Covaris, Woburn, Massachusetts, USA) to obtain DNA fragments of about 350 bp. Accurate profiling of the sonicated sample size distribution was performed on an Agilent 4150 TapeStation System using the High Sensitivity D1000 ScreenTape® (Agilent Technologies, Santa Clara, California, USA). Libraries were prepared using NEBNext® Ultra™ II DNA Library Prep Kit for Illumina® and the NEBNext Multiplex Oligos for Illumina (Dual Index Primers Set 1, New England Biolabs, Ipswich, Massachusetts, USA) and following the manufacturer protocol. In step 3 of the protocol, the size selection of adaptor-ligated DNA was adjusted for each sample according to the average sample size measured with TapeStation. PCR amplification of the adaptor-ligated DNA was performed with eight cycles. Each library was profiled with Qubit and TapeStation assay, as above.

Individual libraries were mixed in 16-sample pools, using equal DNA amounts (200 ng or 250 ng) from samples with similar fragment average sizes. Pool libraries were dried by vacuum centrifugation and resuspended in 7 μl nuclease-free water. Target enrichment was performed using the myBaits Hybridization Capture Kits (Arbor Biosciences, Ann Arbour, MI, USA), following the manufacturer’s protocol v.5.02. A hybridization temperature T_H_ of 60°C was chosen. The temperature T_W_ also corresponded to 60°C. The enriched libraries were amplified with 10 PCR cycles. After amplification, pooled libraries were purified using NucleoSpin™ Gel and PCR Clean-up Kit (Thermo Fisher Scientific, Germany) and characterized by measuring concentration and fragment sizes as mentioned above. The 16-sample pools were again pooled in equimolar quantities and sequenced at the Core Facility Genomics (CF-GEN) of the Helmholtz Zentrum München, Germany (Deutsches Forschungszentrum für Gesundheit und Umwelt, GmbH) on an Illumina Nextseq 1000 Sequencing System.

### Nuclear data analysis

Raw data was first inspected using FastQC v.0.11.8 (Andrews, 2010) and MultiQC v.1.19 (Ewels et al., 2016). PCR duplicates (identical or nearly identical sequences with some mismatches) were removed using ParDRe v.2.2.5, leaving the number of allowed mismatches as the default setting of zero (González-Domínguez and Schmidt, 2016). We used Trimmomatic v.0.39 (Bolger et al., 2014) to cut Illumina adapters and sequences below a quality threshold of 20 and drop the read if below a length of 25bp. Cleaned sequences were inspected again using FastQC and MultiQC to confirm the quality standards. HybPiper v.2.1.5 (Johnson et al., 2016) was used for loci assembly. The best value for SPAdes (Prjibelski et al., 2020) coverage cutoff (--cov_cutoff option) was assigned individually to each sample by running a preliminary assembly with default parameters and calculating the average coverage and standard error per sample using a custom R script. The percent identity threshold for retaining Exonerate hits (--thresh option), and the percentage similarity threshold for the sliding window (--exonerate_hit_sliding_window_thresh option) were both assigned to 85. The option --chimeric_stitched_contig_edit_distance was set to 0, and -- chimeric_stitched_contig_discordant_reads_cutoff was set to 1. Two different target files (option -t_dna) were used to assemble the ingroups (which include only *Astragalus* sequences) and outgroups (which included the other Astragalean species and three outgroup species *Medicago truncatula*, *Trifolium pratense*, and *Cicer arietinum*). Those target sequences consisted of ortholog exons obtained from additional transcriptome data from Su et al. (2021) and other sequences available on NCBI, following the method described in Morales-Briones et al. (2022). We selected the Diamond method (--diamond option) to map reads to the target loci. The function paralog_retriever included in HybPiper was used to recover coding sequences from putative alternative long paralogs. Orthology inference followed Morales-Briones et al. (2022). All scripts used are available at https://bitbucket.org/dfmoralesb/target_enrichment_orthology/src/master/. Sequences were aligned using MACSE v.2.07 (Ranwez et al., 2018). We used Pxclsq v.1.3 (Brown et al., 2017) to remove sites that showed missing data, using a minimum column occupancy of 0.1 (10%). To infer gene trees, we used IQtree v.2.3.0 (Minh et al., 2020). Tips in the gene trees that were mono- and paraphyletic were masked, and the tip with the most unambiguous characters in the trimmed alignment was kept. TreeShrink v.1.3.9 (Mai and Mirarab, 2018) was used to remove abnormally long branches, using a quantile value of 0.1 and excluding outgroups. Homolog fasta files were then generated from those trees and aligned using OMM MACSE v.12.01. To infer orthologs, the monophyletic outgroup (MO) method described by Yang and Smith (2014) was implemented. After aligning and cleaning the ortholog sequences as above, the final ortholog gene trees were reconstructed using IQtree v.2.3.0, using the standard model selection. To produce a quartet-based species tree, ASTRAL v1.19.4.5 was used (Zhang et al., 2020). An aligned supermatrix of fasta sequences was obtained, concatenating all the ortholog gene sequences. This supermatrix was then used to build the concatenated ML species tree using IQtree. To investigate gene tree discordance, PhyParts v.0.0.1 (Smith et al., 2015) and QuartetSampling (QS) v.1.3.1 (Pease et al., 2018) were used. Before running PhyParts, ortholog trees were rooted using *Trifolium pratense*, *Medicago truncatula,* and *Cicer arietinum* as outgroups, and the analysis was executed two times: once with option -a = 1 and -s = 70 (to get the total of concordant and discordant nodes that passes the support threshold of 70%), and once with option -a = 0 (to obtain the number of nodes that did not pass the threshold of 70%). Those two results were combined using an R script and plotted in Python to obtain the proportion of uninformative and missing data. QuartetSampling was executed using the aligned supermatrix as input, with the option -g to include partitions (per loci). Results were plotted with an R script. To investigate potential reticulation relationships in the backbone of *Astragalus*, phylogenetic networks were reconstructed using PhyloNet v.3.8.2 (Wen et al., 2018). For this analysis, 22 taxa representing the 11 major clades retrieved were selected based on the highest number of ortholog sequences, resulting in a dataset that included 576 loci. Sequences of those selected taxa were aligned and clean, and gene trees were built in the same way as described above. Species networks were then inferred using maximum pseudo-likelihood (Yu and Nakhleh, 2015), with the number of reticulations ranging from 0 to 6. The resulting networks and inheritance probabilities were plotted using the Julia package PyPlot (https://github.com/JuliaPy/PyPlot.jl, accessed on 28/08/2024).

### Chloroplast assembly

Off-target chloroplast sequences were assembled using FastPlast v.1.2.9 (McKain and Wilson, 2017). Filtered SPAdes contigs were imported in Geneious v.2023.2.1 (https://www.geneious.com) and mapped to the complete plastome of *A. pattersonii* (NC_063490, ∼140kbp), selecting the option for mapping at high sensitivity. Discordant overlapping contig sequences were manually removed by keeping only the sequence more similar to the reference. A consensus fasta file was then extracted for each sample. Additional complete plastome sequences were downloaded from NCBI (Table S2). A single fasta file, including all the sample sequences, was then produced and aligned using MAFFT v.7.453 (Katoh and Standley, 2013). The alignment was cleaned using Pxclsq, removing columns with more than 40% of the missing data. An ML tree was inferred using IQtree and standard model selection. Abnormally long branching samples were removed from the final tree after visual inspection. Discordance analysis was performed using QuartetSampling. To select the most appropriate model of evolution separately for each plastome region, coding sequences (CDS) were extracted by producing a consensus alignment with annotated *A. pattersonii* complete plastome sequence. ML trees were then reconstructed in IQtree with a partition table for the CDS. Additionally, another tree was reconstructed using a partition table for codon position.

## RESULTS

Target enrichment data for samples representing 80 species were newly generated in this study, of which 77 belong to *Astragalus* and three to other genera in the Astragalean clade (*Colutea*, *Oxytropis*, and *Podlechiella*). The number of raw paired-end reads ranged between 1.2 million and 12 million. The average number of loci with at least 75% length recovery was 701.0, ranging from 484 to 767 (Table S3). There seems to be no correlation between the year of collection of the herbarium specimens (ranging from 1899 to 2014) and the number of loci recovered by HybPiper (Figure S1). HybPiper gave an average of 33.4 paralog warnings per sample, ranging from 3 to 252 (Table S3). On average, 599.8 ortholog sequences per specimen were retrieved, ranging from 230 to 718 (Table S4). The final nuclear data set included 781 MO orthologs with a minimum of 20 taxa per locus. The concatenated matrix resulted in 778,623 bp, with an overall matrix occupancy of 73%. The coalescent-based species tree obtained with ASTRAL (Figure 1) and the concatenated ML tree obtained with IQTree (insert Figure 1, Figure S2) showed very similar topologies and support values (local posterior probability (LLP) and bootstrap (BS), respectively) along the backbone, with some minor differences in the relationships of the deepest nodes. The Astragalean clade and Eu-Astragalus were recovered as monophyletic with high support (LLP = 1, BS = 100), with species placed in clades mostly consistent with previous studies (e.g., Azani et al., 2017; Su et al., 2021; Folk et al., 2024). In Eu-Astragalus, the clades Glottis, Pseudosesbanella, Phaca, Contortuplicata, Hamosa, Trimeniaeus, Incani, Astracantha, Hypoglottis, Diholcos, and Neo-*Astragalus* were recovered with high support except for Astracantha in the coalescent tree (LPP = 0.69, BS = 100) and Hamosa in the concatenated ML tree (LPP = 1, BS = 65). However, unlike Azani et al. (2017), Diholcos was nested in Hypoglottis, splitting the latter into two fully supported clades (LPP = 1, BS = 100). Furthermore, in both our nuclear trees, the Ophiocarpus clade was dispersed and polyphyletic among Hypoglottis taxa (Figures 1, S2), in contrast with our chloroplast dataset, which instead supports a monophyletic Ophiocarpus (BS = 100) sister to Glottis (Figures 2a, S3). Discordance analysis indicates strong monophyly for the Astragalean clade (628 informative genes concordant out of 628, QS score 1/–/0.96) and the Eu-Astragalus + *Oxytropis* (728 informative genes concordant out of 741, QS score 1/–/0.96; Figures 1, S4, S5). However, along the backbone of *Astragalus* and especially for clades in Meso *Astragalus*, high levels of gene tree discordance and high QS skewed frequencies of alternative placement were shown (e.g., 23 informative out of 668 genes with a QS score of 0.27/1/0.48 for the node between Hypoglottis and Diholcos), despite having high support (LLP = 1, BS = 100, Figures 1, S2).

**Figure 1:**
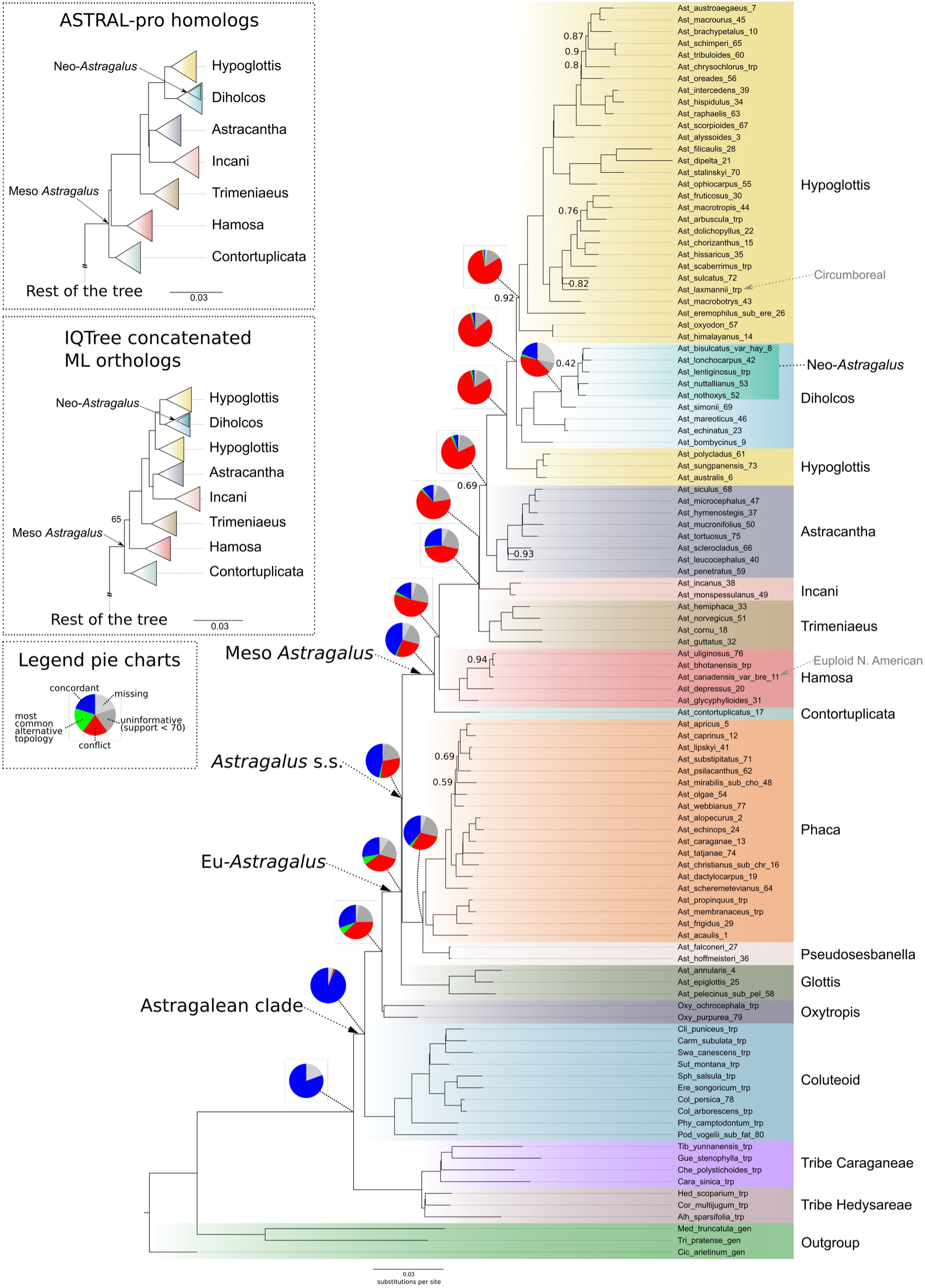
Coalescent-based species tree of Astragalus and few sister clades obtained with ASTRAL and based on 781 ortholog loci. Node support (local posterior probability, LLP) ≥ 0.95 when not indicated. Pie charts indicate gene discordance at nodes along the backbone as calculated by PhyParts. Color coded clade names inside Eu-Astragalus follow Azani et al. (2017) and Su et al. (2021). Inserts shows major differences along the backbone by using different methods as indicated, omitting parts of the trees for which the topology was identical.

**Figure 2:**
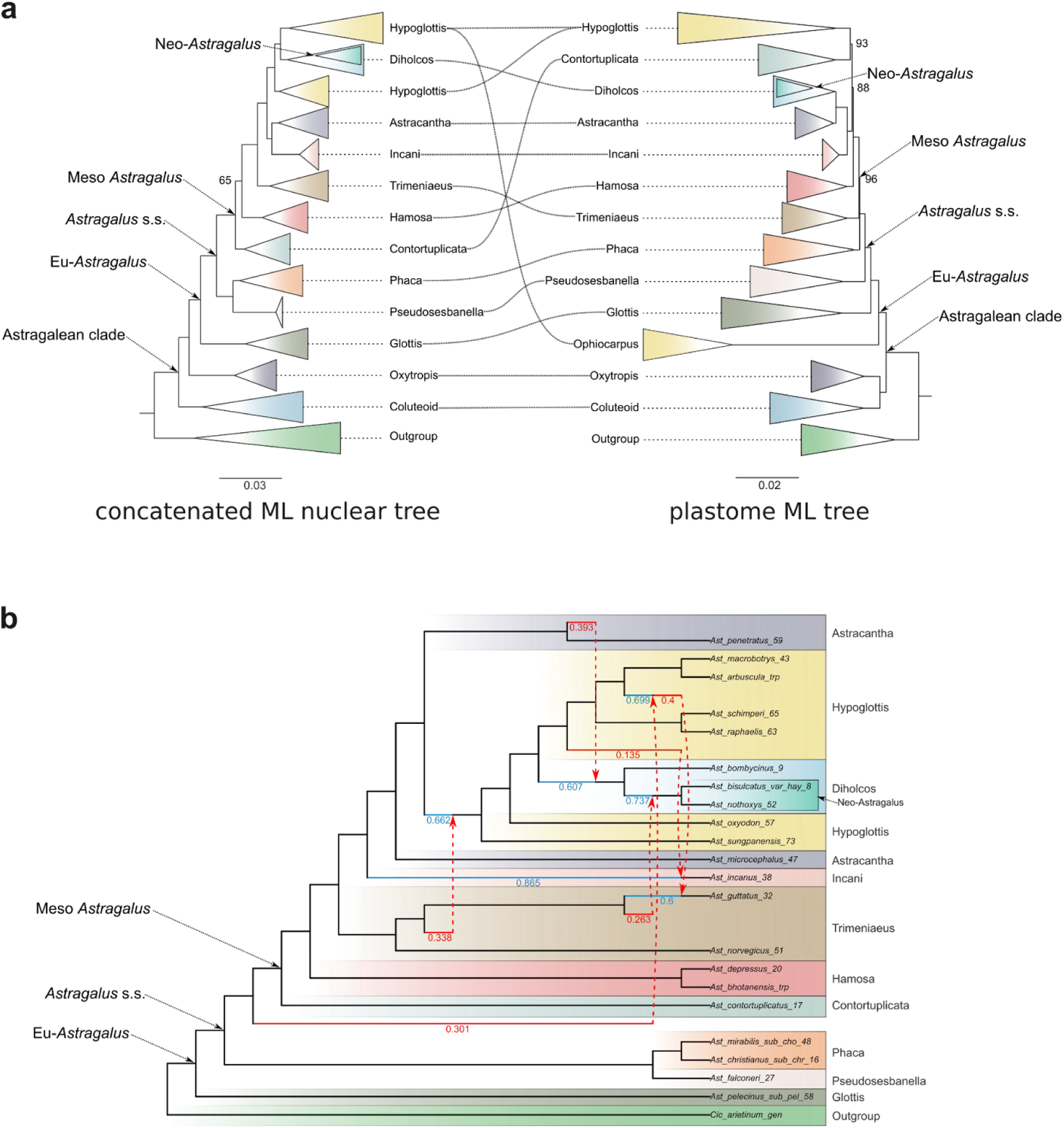
Cytonuclear discordance and nuclear phylogenetic networks suggest presence of hybridization events between Astragalus clades. (a) Comparison of concatenated Maximum Likelihood (ML) nuclear tree (left) and chloroplast ML species tree (right). Bootstrap support = 100 when not indicated. (b) Phylogenetic network obtained with phyloNet with higher probability (6 hybrids, tot log probability = −392572.67). 22 taxa were selected to represent the major clades recovered in the phylogeny.

Phylogenetic networks generated with PhyloNet indicate the presence of extensive reticulate evolution between most of the major *Astragalus* clades (Figure 2b, S6). The most likely number of reticulate events was six based on a log-likelihood of −392572 (Figure S7). The only reticulation event recorded outside Meso *Astragalus* involves a ghost lineage (either not sampled or extinct) sister to Contortuplicata and Hypoglottis clade, with an inheritance probability of 0.3 (Figure 2b). Inside Meso *Astragalus*, several reticulate events with inheritance probability ranging from 0.135 to 0.4 were detected between Trimeniaeus and a common ancestor of Hypoglottis + Diholcos, Trimeniaeus and Neo-Astragalus, Hypoglottis and Incani, Hypoglottis and Trimeniaeus, Astracantha and Diholcos (Fig 2b, S6).

### Chloroplast

Even though chloroplast sequences were not directly targeted, it was possible to obtain chloroplast sequences for all of the samples. The concatenated matrix resulted in 114,580 aligned columns, with 74.6% overall matrix occupancy. There was no appreciable difference between the trees based on plastome data with or without different models of sequence evolution for coding and non-coding sequences. Therefore, subsequent analyses are based on the latter. The resulting ML tree showed high support along the backbone (Figures 2a right, S3), with a topology in agreement with a previous plastome phylogeny (Su et al., 2021). All the clades resolved in our plastome tree, except clade Ophiocarpus, were in agreement with our nuclear trees (both concatenated ML and coalescent-based). Nonetheless, some of them were placed in different positions along the backbone (Figure 2a). Specifically, the Ophiocarpus clade was monophyletic and sister to the Glottis clade in the plastome tree, while it was split in two and nested inside the Hypoglottis clade in the nuclear trees (Figures 1, 2a). Furthermore, Trimeniaeus, Contortuplicata, and Diholcos clades showed different positions inside Eu-Astragalus. The Hypoglottis clade was recovered as monophyletic, with the Contortuplicata clade sister to it in the plastome tree. The latter is sister to Hamosa in the nuclear trees (Figures 1, 2a).

## DISCUSSION

### Capability of bait set to resolve phylogeny

In this study, we produced a robust backbone phylogeny of *Astragalus* by using target enrichment with a clade-specific bait set. Our sample size, with 85 *Astragalus* species, covers about 3% of the Old World and 1% of the New World species. Nevertheless, we selected the representatives carefully to cover all major clades currently supported by molecular data (Azani et al., 2017; Su et al., 2021; Folk et al., 2024). We obtained an overall highly supported species tree, even though we observed large gene tree discordance and cytonuclear discordance (Figures 1, 2a). Our data exhibit low levels of missing or uninformative data (gray and dark gray pie charts Figures 1, S4), excluding low informativeness of loci selected as the main cause of this conflict. Incomplete lineage sorting, often observed in evolutionarily young clades, and reticulate evolution may instead explain better the observed discordances (Smith et al., 2015; Vatanparast et al., 2018; Morales-Briones et al., 2021). Our phylogenetic network analysis supports several events of reticulate evolution between most of the recovered *Astragalus* clades (Figures 2b, S6). Thanks to the large number of loci targeted by our bait set designed specifically for the Astragalean clade, the highly supported backbone of the phylogeny fits, but at the same time, partially challenges the current understanding of the genus, highlighting previously unexplored cytonuclear discrepancies. Our phylogenies recover clades and relationships at unprecedented resolution.

### Overlap with current phylogenies

The recovered nuclear phylogeny shows a strong overlap with current phylogenies in terms of clades recovered, especially with that of Folk et al. (2024; Figure S8). However, contrary to Folk et al. (2024), higher support was obtained, especially within the main clades, and almost all relationships inside the main clades were resolved with high support (LLP = 1). We attribute this to having 5–10 times more sequence data than Folk et al. (2024), allowing for better resolution along the backbone and at shallow nodes.

Similar to Azani et al. (2017), aneuploid Neo-*Astragalus* species were placed as sister to Afro-Mediterranean (and predominantly annual) species in the Diholcos clade, and one euploid American perennial species, *A. canadensis,* was placed close to Asian species in the Hamosa clade (Figure 1). This situation is somewhat similar to Su et al. (2021) which found that the euploid *A. americanus* is nested in the Phaca clade, in agreement with our plastome tree (Figure S3) but not sampled in our nuclear trees. Specifically, aneuploid Neo-*Astragalus* seems to have arisen from annual ancestors of the Diholcos clade, while sampled euploid New World species from the ancestors of Hamosa and Phaca clades, in line with the results of Azani et al. (2017) and Folk et al. (2024) which also found circumboreal species (*A. canadensis* and *A. oreganus*) nesting in the Hamosa clade.

The clade Pseudosesbanella was recovered in agreement with Su et al. (2021) and Folk et al. (2024), though it was resolved with very low support in the latter work. The Glottis clade, recovered with high support and in agreement with Azani et al. (2017) and Su et al. (2021), was represented in Folk et al. (2024) by a single taxon, *A. epiglottis* (syn. *Biserrula epiglottis*; another Glottis member, *A. annularis*, was mentioned in their sample table but it does not appear in their phylogeny), which was placed outside Eu-Astragalus with very low support (<0.5 LPP, Figure S8). Following this result, Folk et al. (2024) supported the segregation of *Biserrula* from *Astragalus*, in disagreement with other studies (e.g., Azani et al., 2017; Su et al., 2021; Wojciechowski, 2005) and ours with strong support (LPP = 1) for the Glottis clade being the earliest diverging clade of Eu-Astragalus. Members of the Glottis clade with a most likely Mediterranean origin (Azani et al., 2019) are annual plants and have very small flowers with a calyx shorter than 4 mm and a standard not exceeding 5 mm (Podlech and Zarre, 2013). The presence of only five fertile stamens in two out of three species forming this clade (i.e., *A. pelecinus* and *A. epiglottis* as well as a unique type of legume that is falcate but strongly dorsi-ventrally compressed/flattened in *A. pelecinus* and *A. biserrula*, are other morphological characteristics that support an isolated position of this clade.

### Cytonuclear discrepancies

We observed some incongruence between nuclear and plastid data (Figure 2a). For example, in the chloroplast tree, the Ophiocarpus clade branches early in *Astragalus*, but according to nuclear data, this group is polyphyletic and nested in the Hypoglottis clade (Figure 2a). Folk et al. (2024) included two species belonging to this clade, which were placed in two different positions: inside the Astracantha and Hypoglottis clades. The position of *A. simonii* in the Diholcos clade and sister to Neo-*Astragalus* in the coalescent-based and concatenated ML nuclear trees is incongruent with previous studies and morphology. Still, in the chloroplast phylogeny, the same sample falls in the Trimeniaeus clade, in agreement with Azani et al. (2017), who used a combination of nuclear and plastid data. Interestingly, when looking at their tree based only on the *matK* gene (Azani et al., 2017, Appendix S6), *A. simonii* was located in the Hypoglottis clade (though with low support). The Ophiocarpus clade also appears monophyletic in the *matK* gene tree but polyphyletic and nested among Diholcos and Hypoglottis taxa in the ITS tree (Azani et al., 2017, Appendix S5). Following our results, *A. stalinskyi* also had an ambiguous position, being placed in Hypoglottis (nuclear data) or Diholcos (chloroplast data – in agreement with Su et al., 2021). Diholcos clade, following nuclear data, was nested inside the Hypoglottis clade (both in coalescent and concatenated ML trees – but see coalescent ASTRAL-pro tree based on homologous sequences, insert in Figure 1), while in the chloroplast phylogeny, it was placed as sister to the Astracantha clade (Figure 2a). Additionally, the clades Contortuplicata and Trimeniaeus were placed in different positions in the two trees.

Similar discrepancies are often observed in plants and are attributed to reticulate evolution (Soltis and Kuzoff, 1995). In *Astragalus*, evidence of reticulate evolution has been reported in the past (Kazemi et al., 2009; Bartha et al., 2013; Záveská et al., 2019; Maylandt et al., 2024). While pinpointing the exact occurrence of hybridization events may be difficult due to the selection of representative taxa, network analysis of the backbone performed in this study supports the presence of several (at least six) reticulate evolutionary events (Figures 2b). This indicates that past hybridization events played an important role in the evolution and establishment of several clades inside the genus. The occurrence of those events mostly in between Meso Astragalus clades overlaps with the high levels of discordance observed along the phylogeny backbone (Figure 1). Thus, ancient hybridization, together with incomplete lineage sorting, may explain the gene tree discordance observed in the genus. Therefore, the current study adds increasing evidence of horizontal gene flow in several *Astragalus* clades.

## CONCLUSIONS

The goal of this study was to build a robust phylogeny of the mega-diverse genus *Astragalus* based on an effective taxon-specific target enrichment bait set. With our carefully selected sampling, we demonstrated that the method successfully obtained a highly supported backbone of the Astragalean clade with great resolution at the subgenus-level relationships in *Astragalus*. Importantly, even with a limited number of sampled taxa, we highlight conspicuous discrepancies between nuclear and plastid signals in *Astragalus*, advocating for caution while using combined loci such as ITS+*matK*. This effort represents only a first step towards a fine-scale resolution of the complex evolutionary history of the lineages in this megagenus. A comprehensive taxon sampling to cover all the morphologically and genetically identified clades and subclades in *Astragalus* is necessary to disentangle intrageneric relationships. Based on current molecular and morphological studies, we anticipate that for a comprehensive phylogeny of the genus in the Old World (∼2600 species), a minimum of about 900 species must be included. These phylogenetic studies will then set the basis for studying the evolution of the largest genus of flowering plants. It will allow us to identify shifts in diversification rate and provide a solid base to study diversification drivers and other important phenomena in a group that impresses for its high diversity.

## Acknowledgments

The authors thank the Elfriede and Franz Jakob Foundation for financial support. The authors thank Katja Arnold and Christina Buhmann for their help with the DNA extractions. SZ acknowledges the support for his morphological and taxonomic studies on the genus *Astragalus* provided by the Iranian National Science Foundation (project number 98029810) and the Alexander von Humboldt Foundation.

## Author contributions

DM, DB, and GK conceived the idea. DB and DM performed the experiments and conducted formal analysis. SZ and AL contributed with species identifications and critical revisions. All aouthors contributed to the final draft of the manuscript.

## Data availabity

Target enrichment data generated for this study can be found in the NCBI BioProject XXXXX (please refer to Table XXX for SRA accession numbers). Analyses files are available from the Dryad repository https://doi.org/10.5061/dryad.XXXX

---

**Figure S1:**
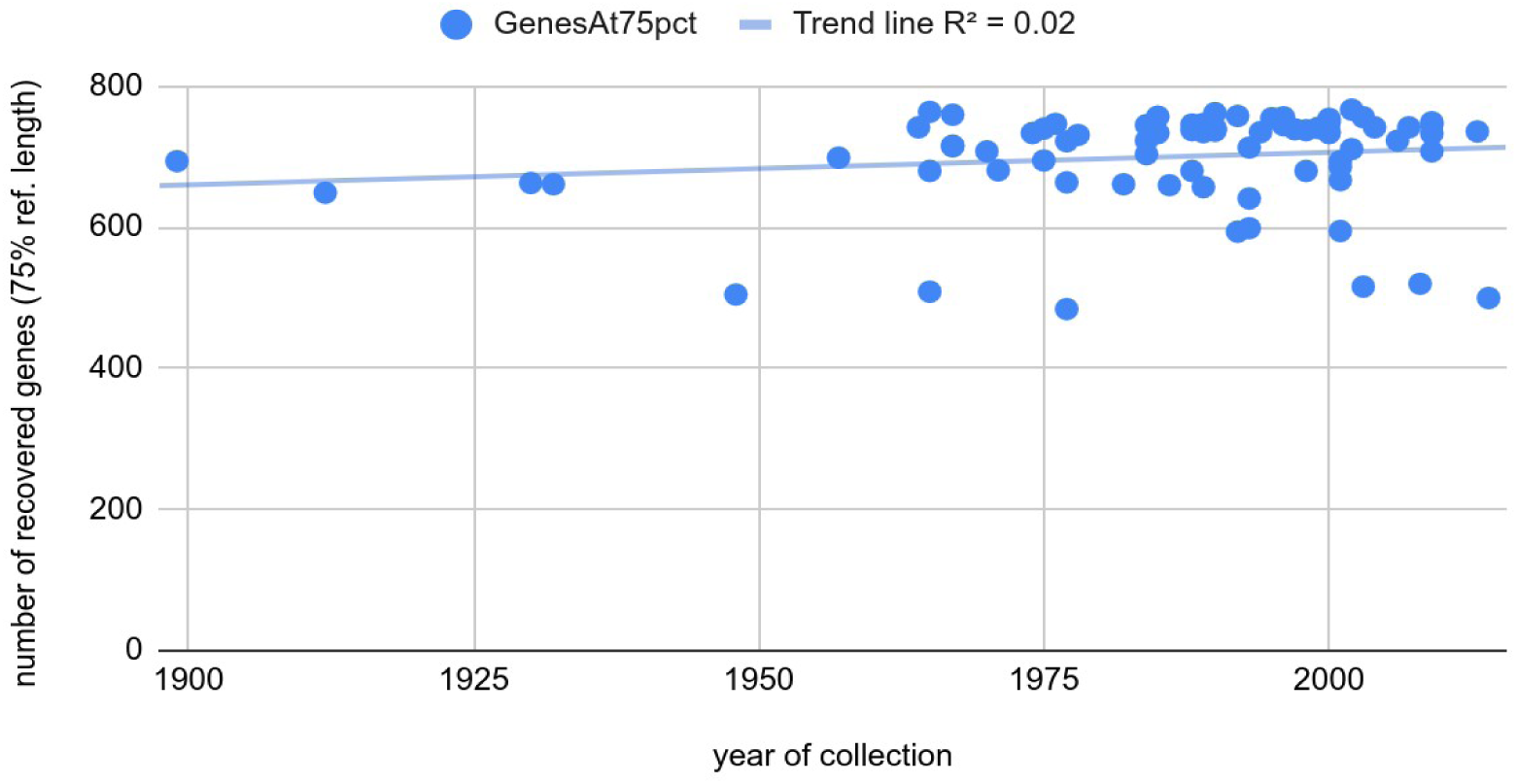
Number of recovered genes (with at least 75% percent of reference sequence length) vs age of herbarium specimen. No clear correlation was observed (trend line R2 = 0.02).

**Figure S2:**
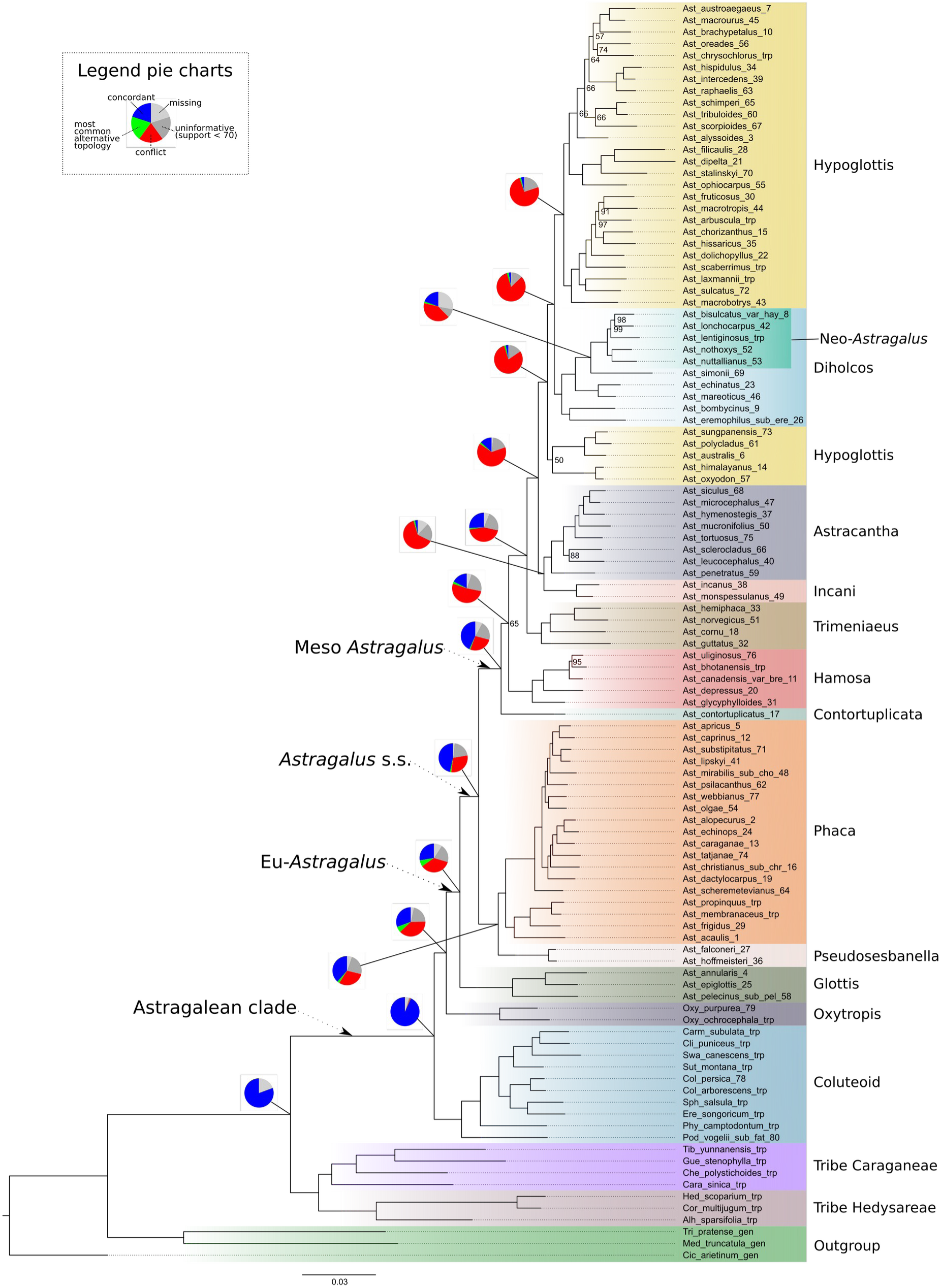
Concatenated maximum likelihood tree (IQtree) produced by concatenating all sequences in a single supermatrix with a total length of 778,623bp and 27% missing data. Bootstrap support on nodes = 100 when not indicated. Pie charts indicate gene discordance at nodes along the backbone calculated with PhyParts. Colour-coded clade names inside Eu-Astragalus matches Azani et al. (2017) and Su et al. (2021).

**Figure S3.**
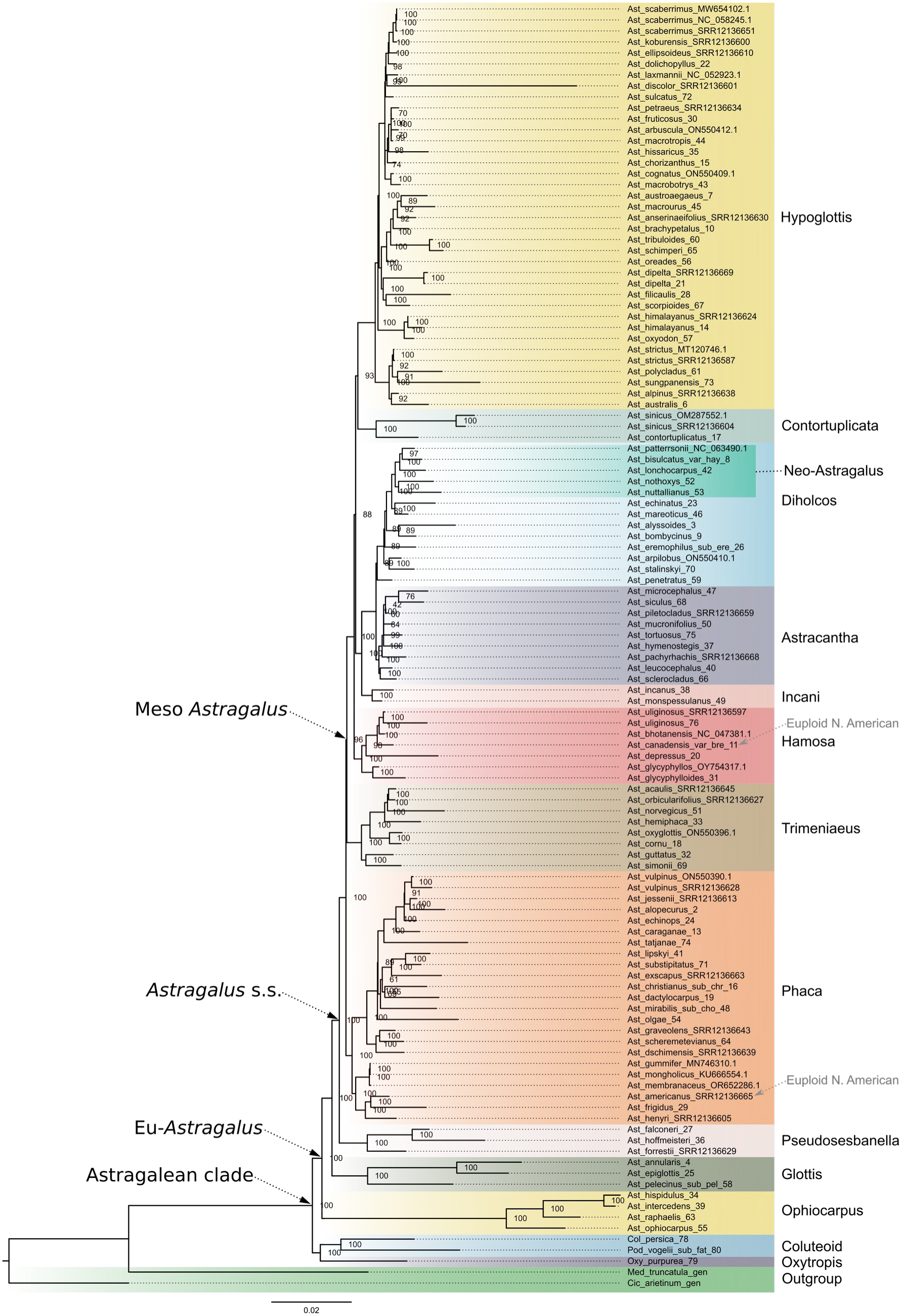
Maximum likelihood (IQtree) species tree inferred from plastome sequences, based on 114,580 aligned columns with 74.6% overall matrix occupancy. Numbers beside nodes indicate bootstrap support. Colour-coded clade names inside Eu-Astragalus matches Azani et al. (2017) and Su et al. (2021).

**Figure S4:**
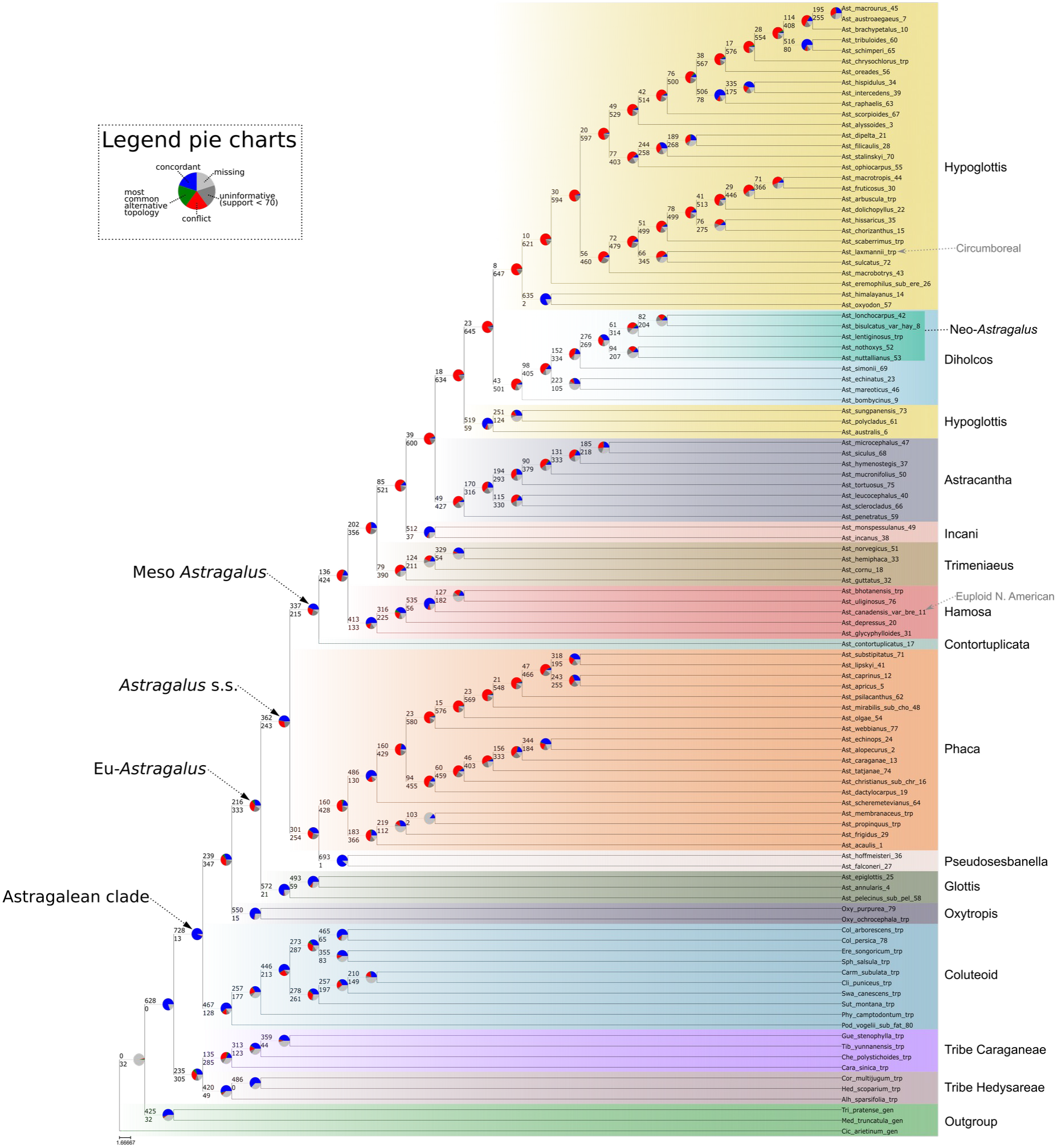
PhyParts analysis results mapped on the coalescent-based species tree obtained with ASTRAL. Uninformative values are ones with bootstrap support < 70.

**Figure S5:**
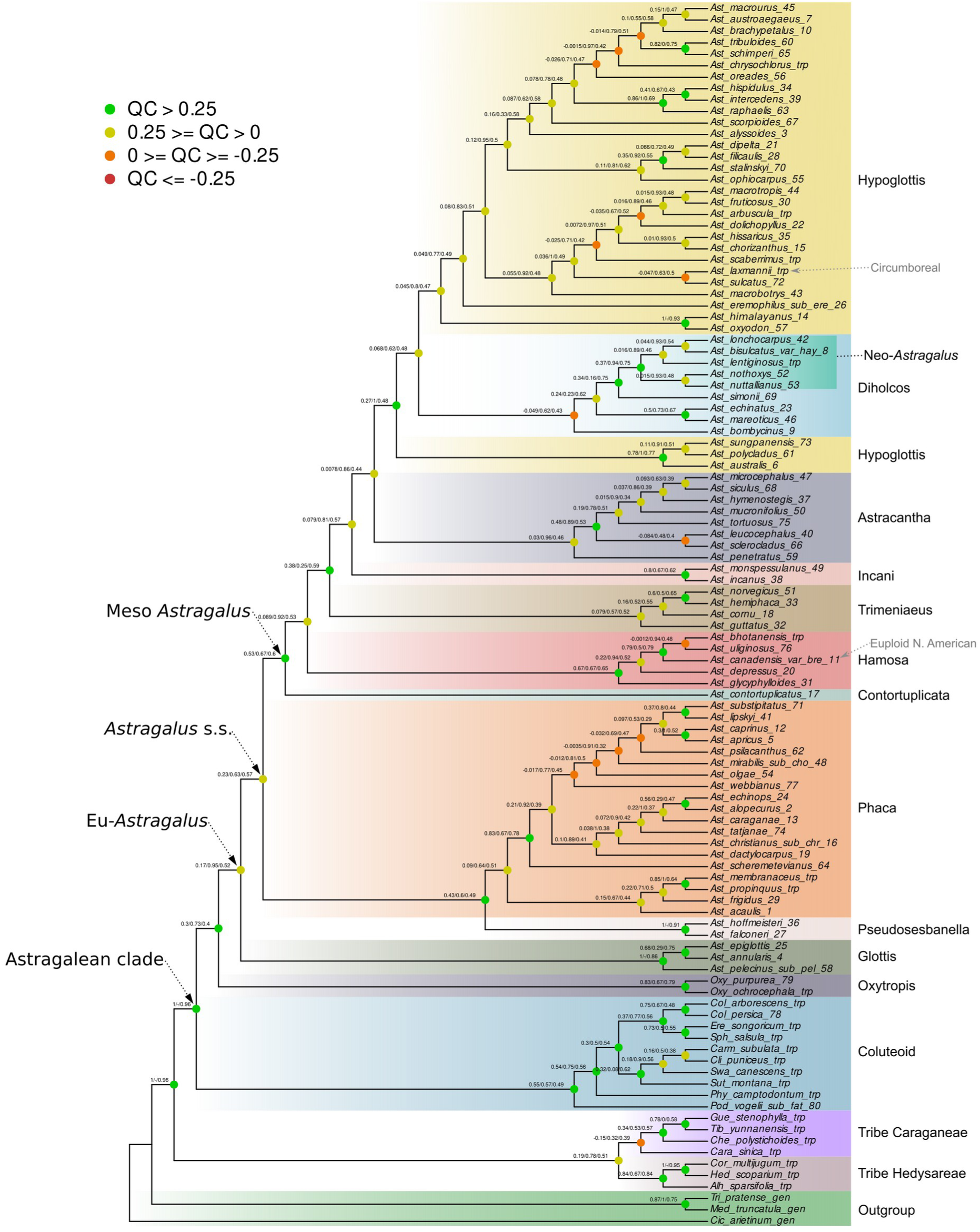
Quartet Sampling probabilities mapped on the coalescent-based species tree obtained with ASTRAL.

**Figure S6:**
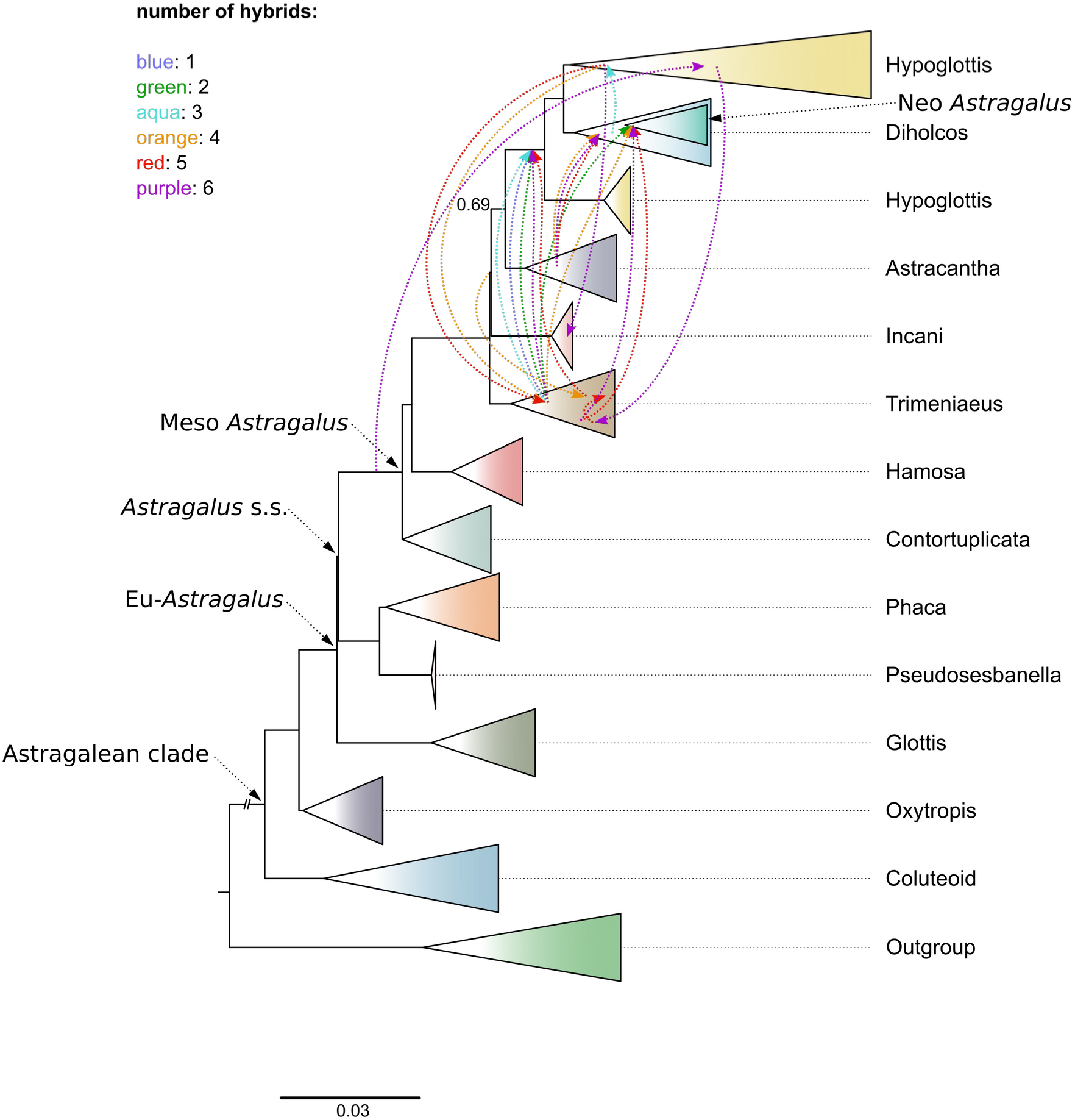
Approximation of reticulate events among clades detected with PhyloNet analysis overlapped on Astral phylogeny (Fig. 1) setting the number of hypothesized hybrids from 1 to 6.

**Figure S7:**
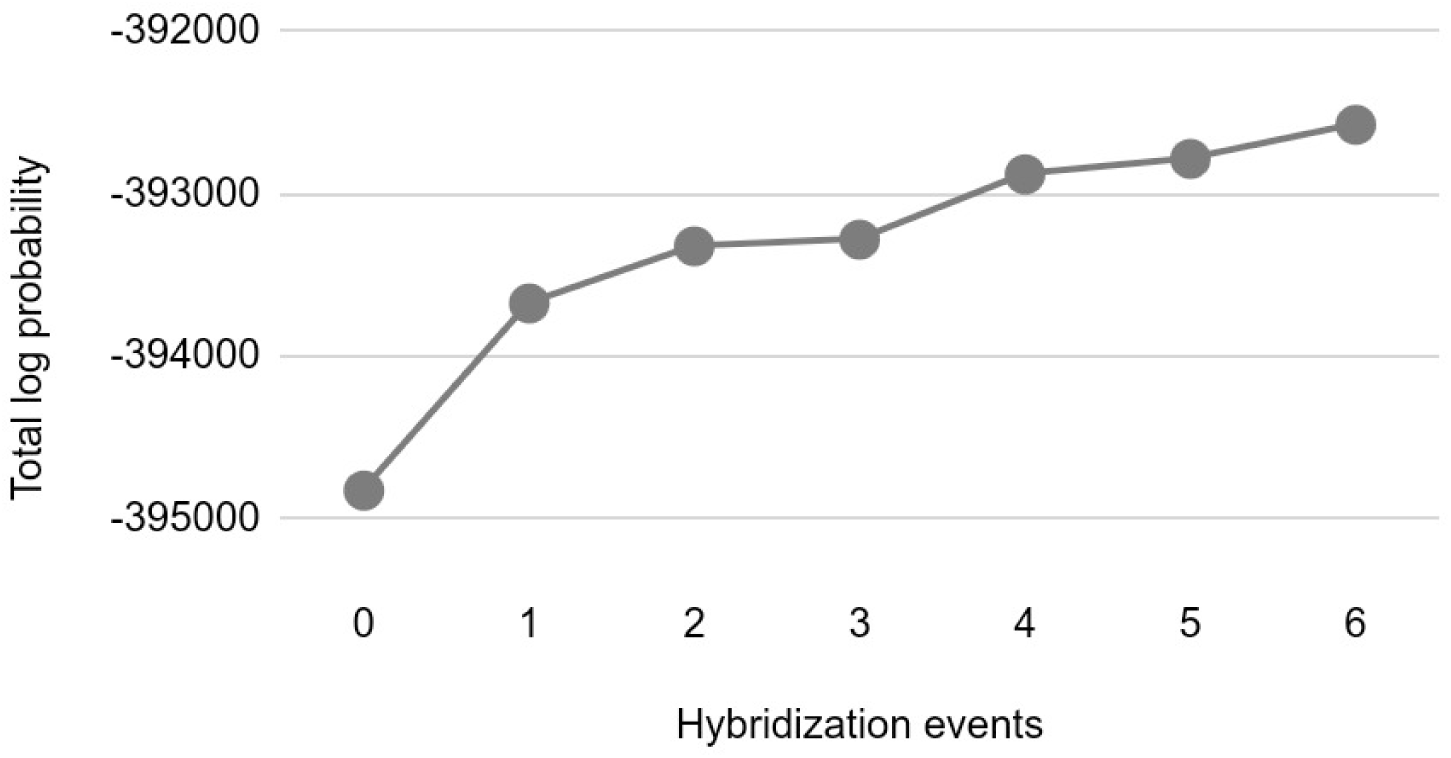
Total log probability vs number of hybridization events inferred by PhyloNet analysis based on 22 taxa (576 loci). The most likely number of reticulate events was six based on a log-likelihood of −392572.

**Figure S8:**
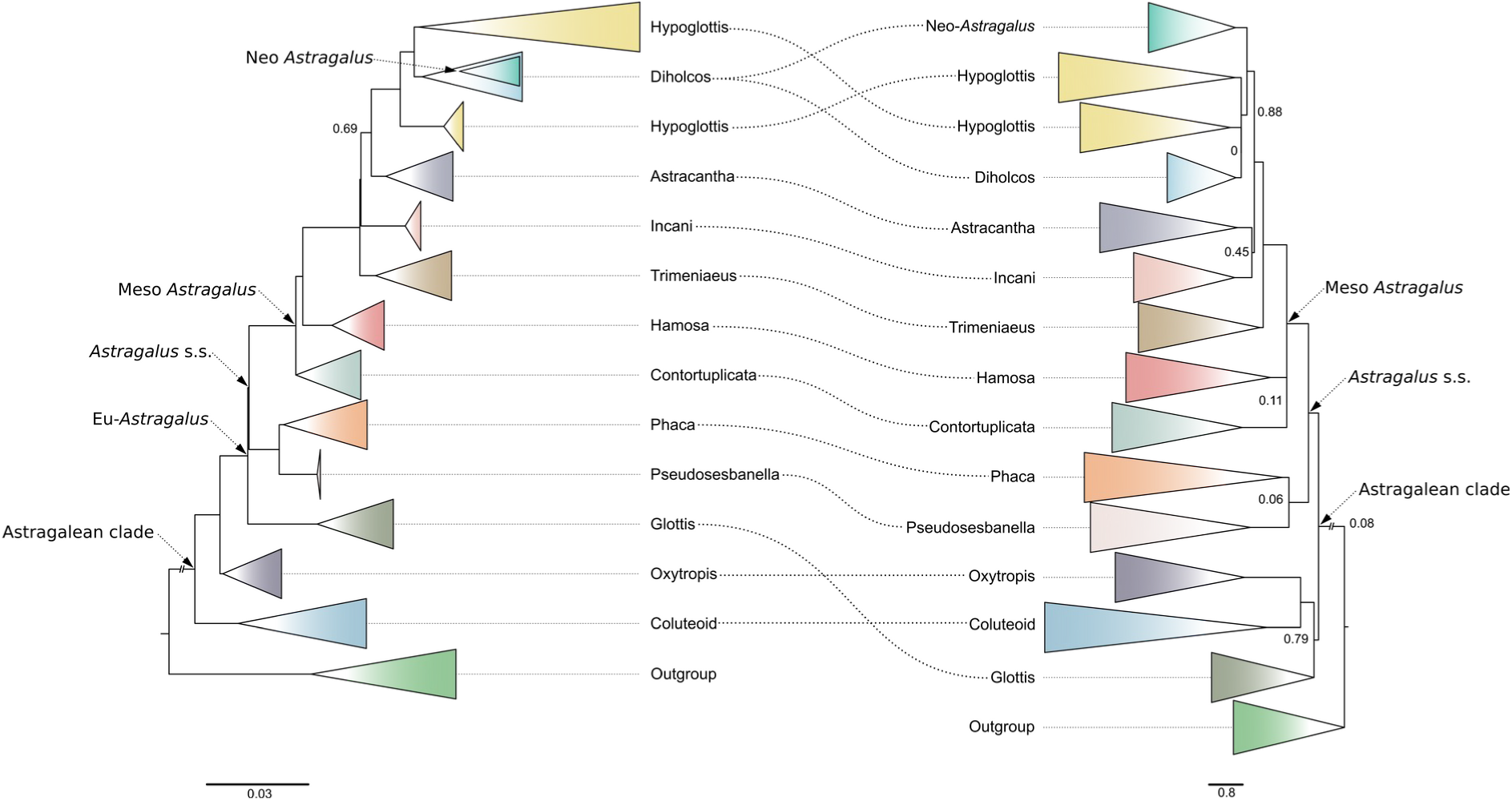
Comparison between coalescent-based species tree (ASTRAL) produced in this study (left) and the one produced by Folk et al 2024 (right). Node support (local posterior probability) ≥ 0.95 when not indicated.

**Table S1:** Taxa sampled in Azani et al. 2017 which were resampled in this study. In grey, taxa for which a different voucher was used.

| No. | Taxon | Clade sensu Anzani et al 2017 | Section | Voucher Azani et al 2017 | Voucher Sampled | Origin | year of collection |
| --- | --- | --- | --- | --- | --- | --- | --- |
| 7 | *A. alyssoides* Lam. | Hypoglottis | Hololeuce | A. Talebi, 43138, TUH | Safavi & Alizade & Nikchehre, 84119, 110283 MSB | Iran | 2003 |
| 9 | *A. annularis* Forssk. | Glottis |  | K. Tilbörger, s.n., MSB | K. Tilbörger, s.n., MSB | Israel | 1992 |
| 21 | *A. bisulcatus* A. Gray var. *haydenianus* | Diholcos | Bisulcati | M. F. Wojciechowski, 133 | A. Tiehm, 13563, M | USA | 2001 |
| 23 | *A. bombycinus* Boiss. | Diholcos | Platylglottis | K. H. Rechinger, 9857, M | Rechinger, 9857, M | Iraq | 1957 |
| 26 | *A. brachypetalus* Trautv. | Hypoglottis | Stereothrix | H. Akhani, 9390, M | H. Akhani, 9390, M | Iran | 1994 |
| 35 | *A. canadensis* L. var. *brevidens* | Hamosa | Uliginosi | M. F. Wojciechowski, 302 | A. Tiehm, 13225, 86946 M | Canada | 2000 |
| 37 | *A. caprinus* L. subsp. *caprinus* | Phaca | Caprini | O. Andres, 2774, MSB | O. Andres, 2774, 5932 MSB | Iraq | 1969 |
| 39 | *A. christianus* L. subsp. *christianus* | Phaca | Astragalus | M. Nydegger, 47315, MSB | M. Nydegger, 47424, 11057, MSB | Turkey | 1993 |
| 44 | *A. contortuplicatus* L. | Contortuplicata | Cycloglottis | Botschkin and Klinkova, 15188, MSB | Botschkin and Klinkova, 15188, 5882 MSB | Russia-Europe | 1989 |
| 45 | *A. cornu-bovis* Lipsky | Trimeniaeus | Sesamei | K. H. Rechinger, 51405, M | K. H. Rechinger, 51405, 257255 M | Iran | 1975 |
| 50 | *A. depressus* L. | Hamosa | Tapinodes | W. Lang, s.n., M | W. Lang, s.n., M | Switzerland | 1988 |
| 51 | *A. dipelta* Bunge | Hypoglottis | Dipelta | F. Memariani and Zangoeei, 42542, FUMH | Assadi & Maassoumi, 50863, 6986 M | Iran | 1984 |
| 53 | *A. echinatus* Murr. | Diholcos | Pentaglottis | D. Podlech, 51232, MSB | D. Podlech, 51232, 002549 MSB | Spain | 1996 |
| 55 | *A. epiglottis* L. (1) | Glottis | Epiglottis | D. Podlech, 45878, MSB | D. Podlech, 45878, 7157 MSB | Morocco | 1989 |
| 58 | *A. eremophilus* Boiss. subsp. *eremophilus* (1) | Diholcos | Harpilobus | Soják, 1534, MSB | Maassoumi & Abouhamzeh, 52074, 005978 MSB | Iran | 1985 |
| 64 | *A. filicaulis* (2) | Hypoglottis | Sesamei | M. R. Joharchi, 34074, FUMH | Assadi & Maassoumi, 55373, 7540 MSB | Iran | 1986 |

| No. | Taxon | Clade sensu Anzani et al 2017 | Section | Voucher Azani et al 2017 | Voucher Sampled | Origin | year of collection |
| --- | --- | --- | --- | --- | --- | --- | --- |
| 67 | *A. flavus* Nutt. | Diholcos | Ocreati | M. F. Wojciechowski, 138, W | O. Buchtien, s.n., M | USA | 1895 |
| 72 | *A. glycyphyllodes* DC. | Hamosa | Glycyphyllus | A. Ghahreman and V. Mozaffarian, 9661, TUH | A. Ghahreman and V. Mozaffarian, 9661, 100537 MSB | Iran | 1990 |
| 75 | *A. guttatus* Banks & Sol. (1) | Trimeniaeus | Aulacolobus | Sytin and Ryazanova, s. n., MSB | Sytin, 002777 MSB | Ukraine | 2000 |
| 79 | *A. hemiphaca* Kar. & Kir. | Trimeniaeus | Oroboidei | Saposhnikov and Schischkin, s.n., M | [cyrillic], 5420 M | Kazakhstan | 1912 |
| 80 | *A. himalayanus* Klotzsch | Hypoglottis | Komaroviella | Dickoré, 20743, MSB | Dickoré, 20743, 157810 MSB | India | 2005 |
| 81 | *A. hispidulus* DC. | Ophiocarpus | Hispiduli | D. Podlech, 44568, MSB | D. Podlech, 44568, 005977 MSB | Egypt | 1989 |
| 82 | *A. hissaricus* Lipsky | Hypoglottis | Hololeuce | Varivtseva and Nepli, 517, MSB | Varivtseva & Nepli, 517, 71531 MSB | Tajikistan | 1948 |
| 83 | *A. hymenostegis* Fisch. & C.A.Mey. | Astracantha | Hymenostegis | Y. Salmaki et al., 39869, TUH | Bagheri, 98071, 168221 MSB | Azerbaijan | 2013 |
| 85 | *A. incanus* L. subsp. *incanus* | Cercidothrix | Inceni | Lambinon, 85/Es/234, MSB | Lambinon, 139789 MSB | Spain | 1985 |
| 87 | *A. intercedens* Sam. ex Rech.f. | Ophiocarpus | Hispiduli | Colonette, 6513, MSB | Colonette, 6513, 8086 MSB | Saudi Arabia | 1988 |
| 92 | *A. latifolius* Lam. | Cercidothrix | Inceni | S. Zarre et al., 3910, TUH | Safavi & Alizade & Nikchehre, 84136, 110287 MSB | Iran | 2003 |
| 95 | *A. leucocephalus* Graham | Ophiocarpus | Poliiothrix | Schickhoff, 260, MSB | Schickhoff, 260, 142142 MSB | Pakistan | 1990 |
| 97 | *A. lonchocarpus* Torr. | Diholcos | Lonchocarpi | M. F. Wojciechowski, 143 | N.H. Holmegren, 11924, M | USA | 1993 |
| 103 | *A. macrourus* Fisch. & C.A.Mey. | Hypoglottis | Malacothrix | D. Podlech and A. A. Maassoumi, 55251, MSB | Podlech & Maassoumi & Zarre, 55251, 001838 MSB | Iran | 2001 |
| 104 | *A. mareoticus* Delile | Diholcos | Harpilobus | D. Podlech, 49326, MSB | D. Podlech, 49256, 9466 MSB | Morocco | 1990 |
| 106 | *A. microcephalus* Willd. | Astracantha | Rhacophorus | D. Podlech, A. A. Maassoumi and S. Zarre, 55225, MSB | Podlech, 55200, 104726 MSB | Iran | 2001 |

| No. | Taxon | Clade sensu Anzani et al 2017 | Section | Voucher Azani et al 2017 | Voucher Sampled | Origin | year of collection |
| --- | --- | --- | --- | --- | --- | --- | --- |
| 109 | *A. norvegicus* Weber | Trimeniaeus | Oroboidei | Buttler and Gauhl, 8260, M | Buttler and Gauhl, 8260, M | Norway | 1965 |
| 110 | *A. nothoxys* A. Gray | Diholcos | Leptocarpi | M. F. Wojciechowski, 177 | A. Meebold, 15315 M | USA | 1932 |
| 111 | *A. nuttallianus* DC. | Diholcos | Leptocarpi | M. F. Wojciechowski, 101 | A. Meebold, 8820, M | USA - Arizona | 1930 |
| 114 | *A. ophiocarpus* Bunge | Ophiocarpus | Ophiocarpus | Faghihnia and Zangoie, 30133, FUMH | D. Podlech, 30832, 10892 MSB | Iran | 1978 |
| 116 | *A. oreades* C.A.Mey. | Hypoglottis | Hypoglottidei | Ekici, 3768, MSB | Ekici, 9768, 146896 MSB | Turkey | 2007 |
| 119 | *A. oxyodon* Baker | Hypoglottis | Komaroviella | Millinger, 3509, MSB | A. Millinger, 3509, 74948, MSB | Pakistan | 1999 |
| 122 | *A. pelecinus* subsp. *pelecinus* | Glottis | Biserrula | D. Podlech, 51802, MSB | D. Podlech, 51802, 005973 MSB | Spain | 1993 |
| 123 | *A. penetratus* Maassoumi |  | Brachylobium | Parishani, 14197, M | Parishani, 14197, M | Iran | 2003 |
| 126 | *A. persepolitanus* Boiss. | Trimeniaeus | Sesamei | S. Zarre and N. Azani, 43861, TUH | Zarre & Isalkmaki & Ebrahimi, 34339, 155256 MSB | Iran | 2009 |
| 128 | *A. polycladus* Bureau & Franch. | Hypoglottis | Poliiothrix | Dickoré, 14096, MSB | Dickoré, 14096, 70343 MSB | China | 1996 |
| 131 | *A. raphaelis* G.Ferro | Ophiocarpus | Sesamei | Brullo, Giusso and Sciandrello, s.n., MSB | Brullo & Giusso & Sciandrello, 138169 MSB | Italy | 2006 |
| 135 | *A. schimperi* Boiss. | Hypoglottis | Sesamei | A. Danin, s.n., MSB | A. Danin, 10033, MSB | Israel | 1988 |
| 137 | *A. sclerocladus* Bunge | Astracantha | Acanthophace | American-Iranian Botanical Delegation, 33731, MSB | American-Iranian Botanical Delegation, 33731, 116432 MSB | Iran | 2004 |
| 138 | *A. scorpioides* Pourr. ex Willd. | Hypoglottis | Sesamei | R. Vogt and Ch. Oberprieler, 8802, 14493 MSB | Vogt & Oberprieler, 18-427, 005965 MSB | Morocco | 1995 |
| 145 | *A. simonii* Hub.-Mor. | Trimeniaeus | Theiochrus | M. Nydegger, 46240, MSB | Nydegger, 46239, 8634 MSB | Turkey | 1992 |
| 150 | *A. stalinskyi* Širj. | Diholcos | Ankylotus | F. Memariani and Zangoie, 42341, FUMH | Assadi & Maassoumi, 50866, 8693 MSB | Iran | 1984 |
| 166 | *A. uliginosus* L. | Hamosa | Uliginosi | Korotkova and Kovaleva, 993, MSB | Korotkova & Kovaleva, 993, 92743 MSB | Russia-Siberia | 1977 |

| No. | Taxon | Clade sensu Anzani et al 2017 | Section | Voucher Azani et al 2017 | Voucher Sampled | Origin | year of collection |
| --- | --- | --- | --- | --- | --- | --- | --- |
| 172 | *Colutea persica* Boiss. | Outgroup |  | Manutscgehri, 970, MSB | D. Podlech, 970, 75414 MSB | Iran | 1977 |
| 175 | *Oxytropis purpurea* Markgr. | Outgroup | - | A.O. Chater, 303, M | A.O. Chater, 303, M | Yugoslavia | 1971 |
| 178 | *Podlechiella vogelii* subsp. *fatimensis* | Outgroup | - | D. Podlech, 35852, MSB | D. Podlech, 36700, MSB | Algeria | 1982 |

**Table S2.**
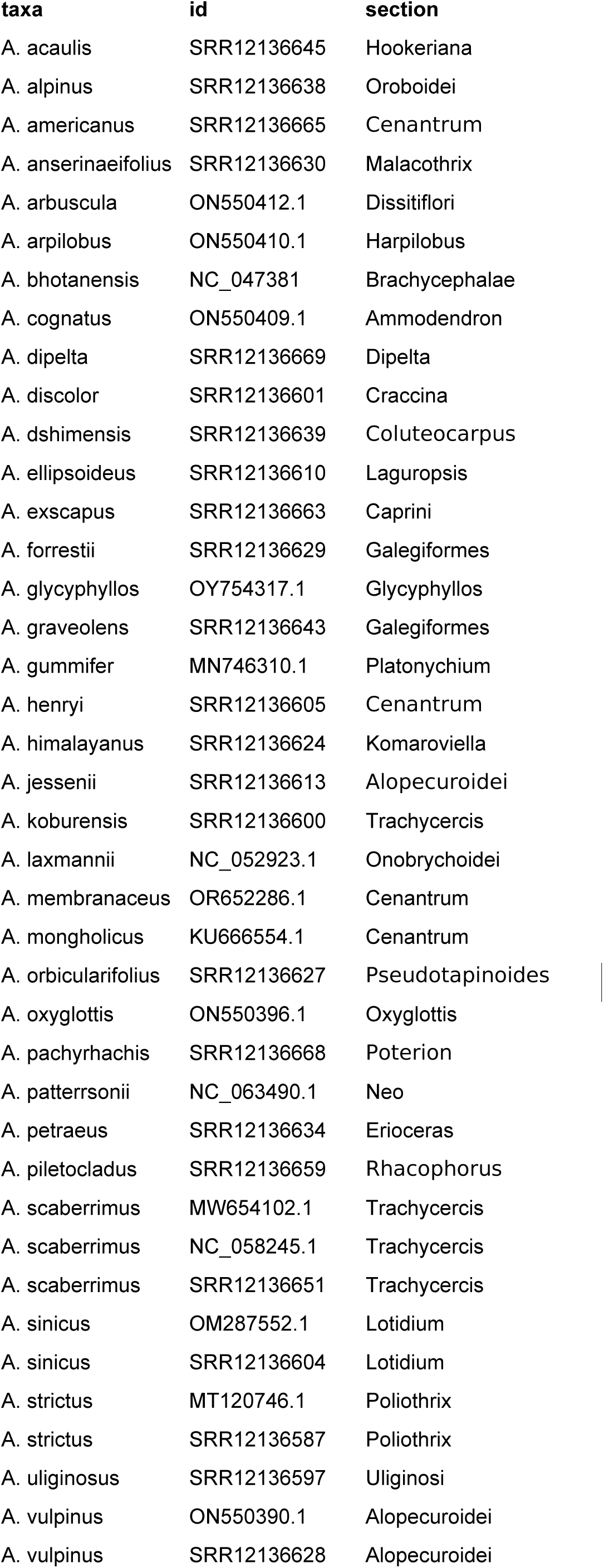
Additional plastome sequences downloaded from NCBI database used in this study.

| **taxa** | **id** | **section** |
| --- | --- | --- |
| *A. acaulis* | SRR12136645 | Hookeriana |
| *A. alpinus* | SRR12136638 | Oroboidei |
| *A. americanus* | SRR12136665 | Cenantrum |
| *A. anserinaefolius* | SRR12136630 | Malacothrix |
| *A. arbuscula* | ON550412.1 | Dissitiflori |
| *A. arpilobus* | ON550410.1 | Harpilobus |
| *A. bhotanensis* | NC_047381 | Brachycephalae |
| *A. cognatus* | ON550409.1 | Ammodendron |
| *A. dipelta* | SRR12136669 | Dipelta |
| *A. discolor* | SRR12136601 | Craccina |
| *A. dshimensis* | SRR12136639 | Coluteocarpus |
| *A. ellipsoideus* | SRR12136610 | Laguropsis |
| *A. exscapus* | SRR12136663 | Caprini |
| *A. forrestii* | SRR12136629 | Galegiformes |
| *A. glycyphyllos* | OY754317.1 | Glycyphyllos |
| *A. graveolens* | SRR12136643 | Galegiformes |
| *A. gummifer* | MN746310.1 | Platonychium |
| *A. henryi* | SRR12136605 | Cenantrum |
| *A. himalayanus* | SRR12136624 | Komaroviella |
| *A. jessenii* | SRR12136613 | Alopecuroidei |
| *A. koburensis* | SRR12136600 | Trachycercis |
| *A. laxmannii* | NC_052923.1 | Onobrychoidei |
| *A. membranaceus* | OR652286.1 | Cenantrum |
| *A. mongholicus* | KU666554.1 | Cenantrum |
| *A. orbicularifolius* | SRR12136627 | Pseudotapinoidea |
| *A. oxyglottis* | ON550396.1 | Oxyglottis |
| *A. pachyrhachis* | SRR12136668 | Poterion |
| *A. pattersonii* | NC_063490.1 | Neo |
| *A. petraeus* | SRR12136634 | Erioceras |
| *A. piletocladus* | SRR12136659 | Rhacophorus |
| *A. scaberrimus* | MW654102.1 | Trachycercis |
| *A. scaberrimus* | NC_058245.1 | Trachycercis |
| *A. scaberrimus* | SRR12136651 | Trachycercis |
| *A. sinicus* | OM287552.1 | Lotidium |
| *A. sinicus* | SRR12136604 | Lotidium |
| *A. strictus* | MT120746.1 | Poliiothrix |
| *A. strictus* | SRR12136587 | Poliiothrix |
| *A. uliginosus* | SRR12136597 | Uliginosi |
| *A. vulpinus* | ON550390.1 | Alopecuroidei |
| *A. vulpinus* | SRR12136628 | Alopecuroidei |

**Table S3:**
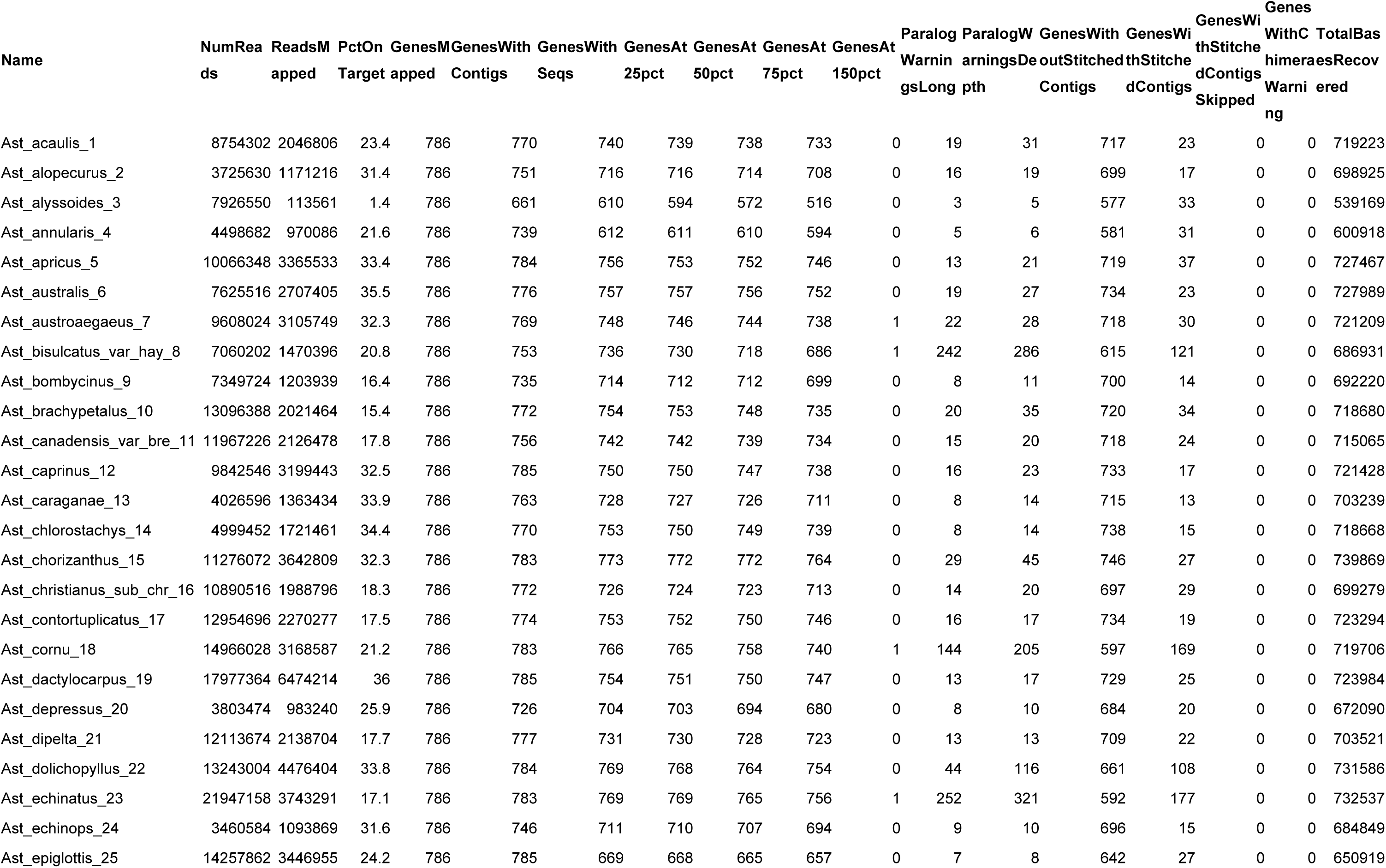
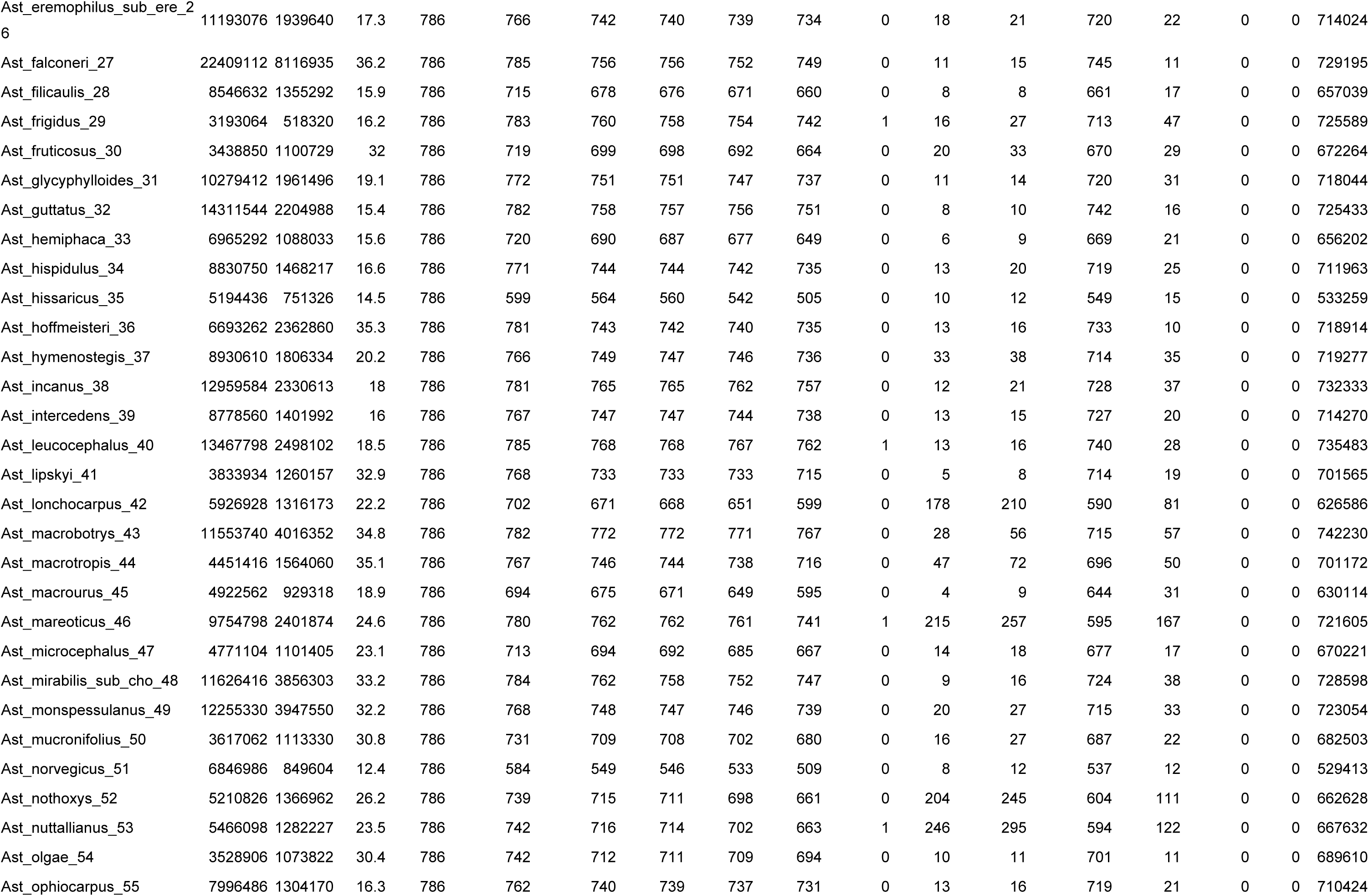
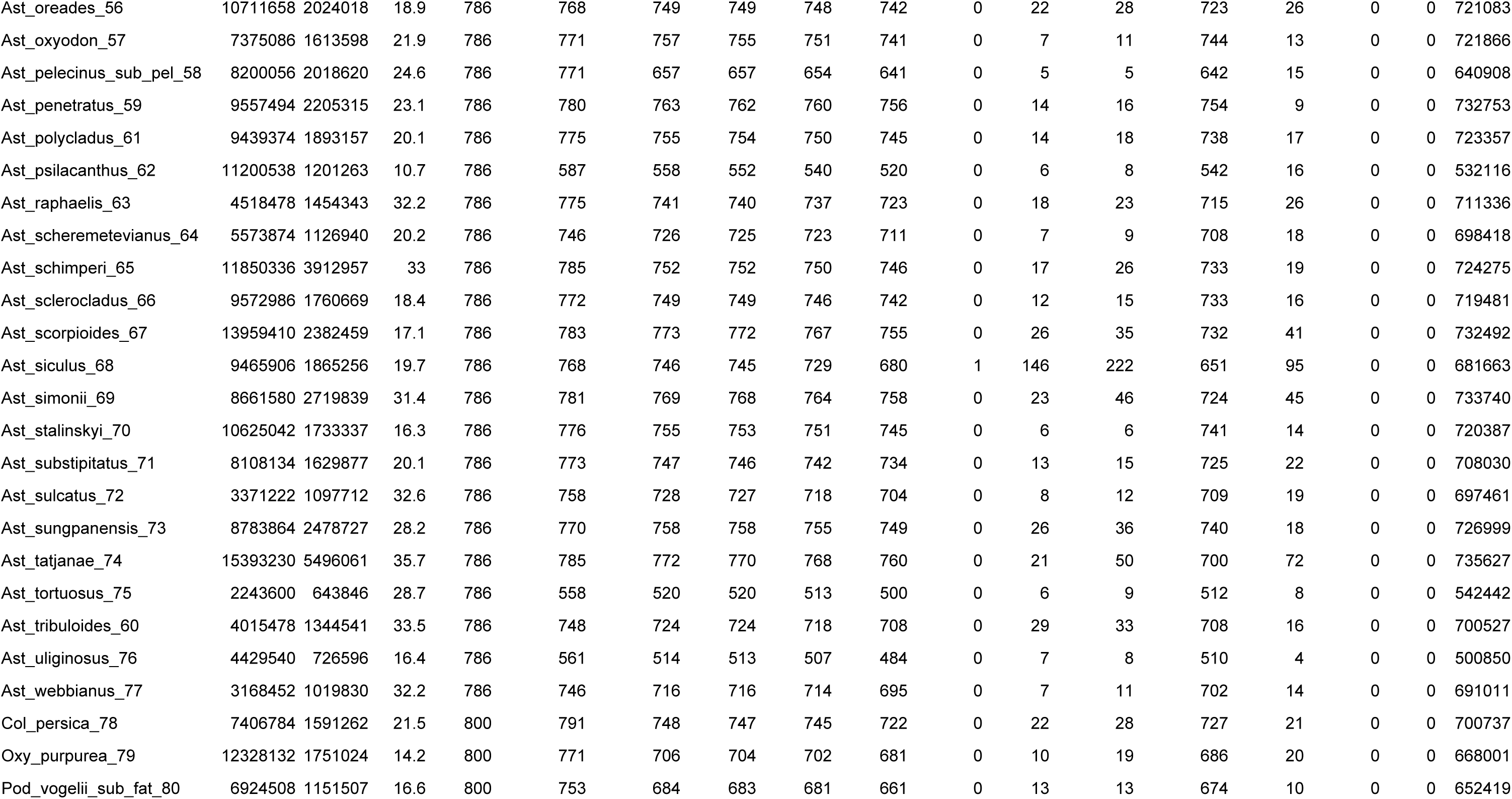
Hybpiper gene recovery statistics.

**Table S4:**
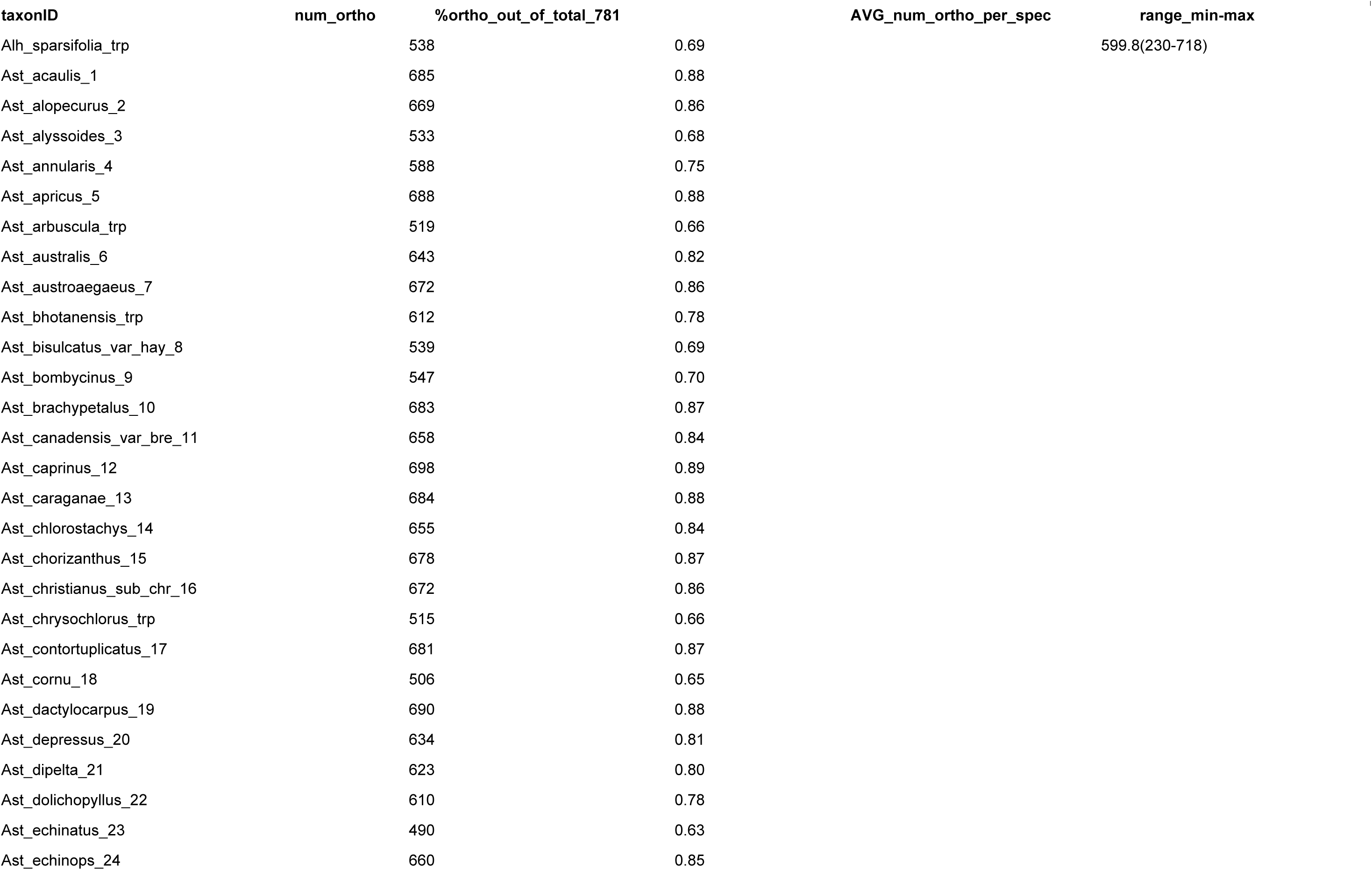
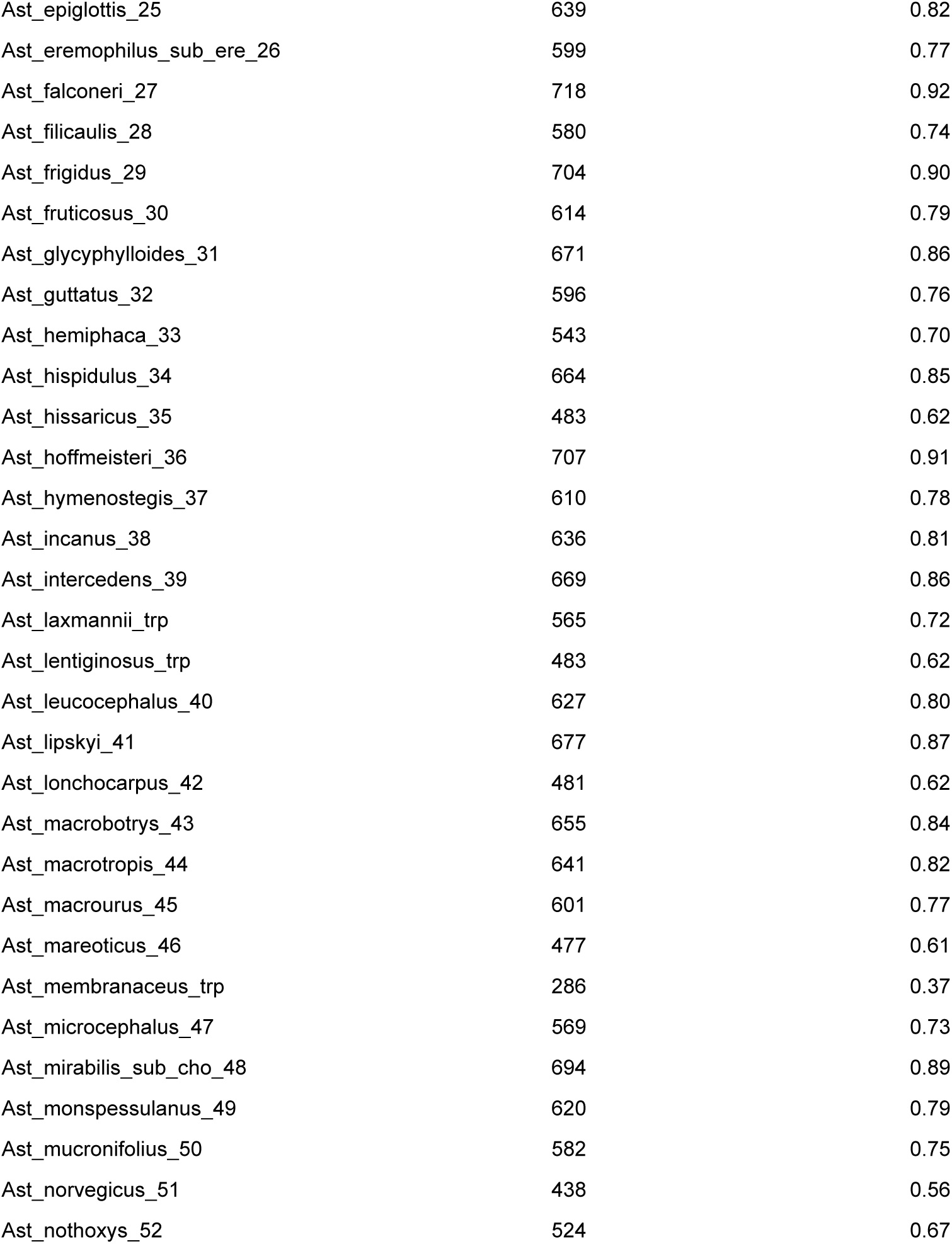
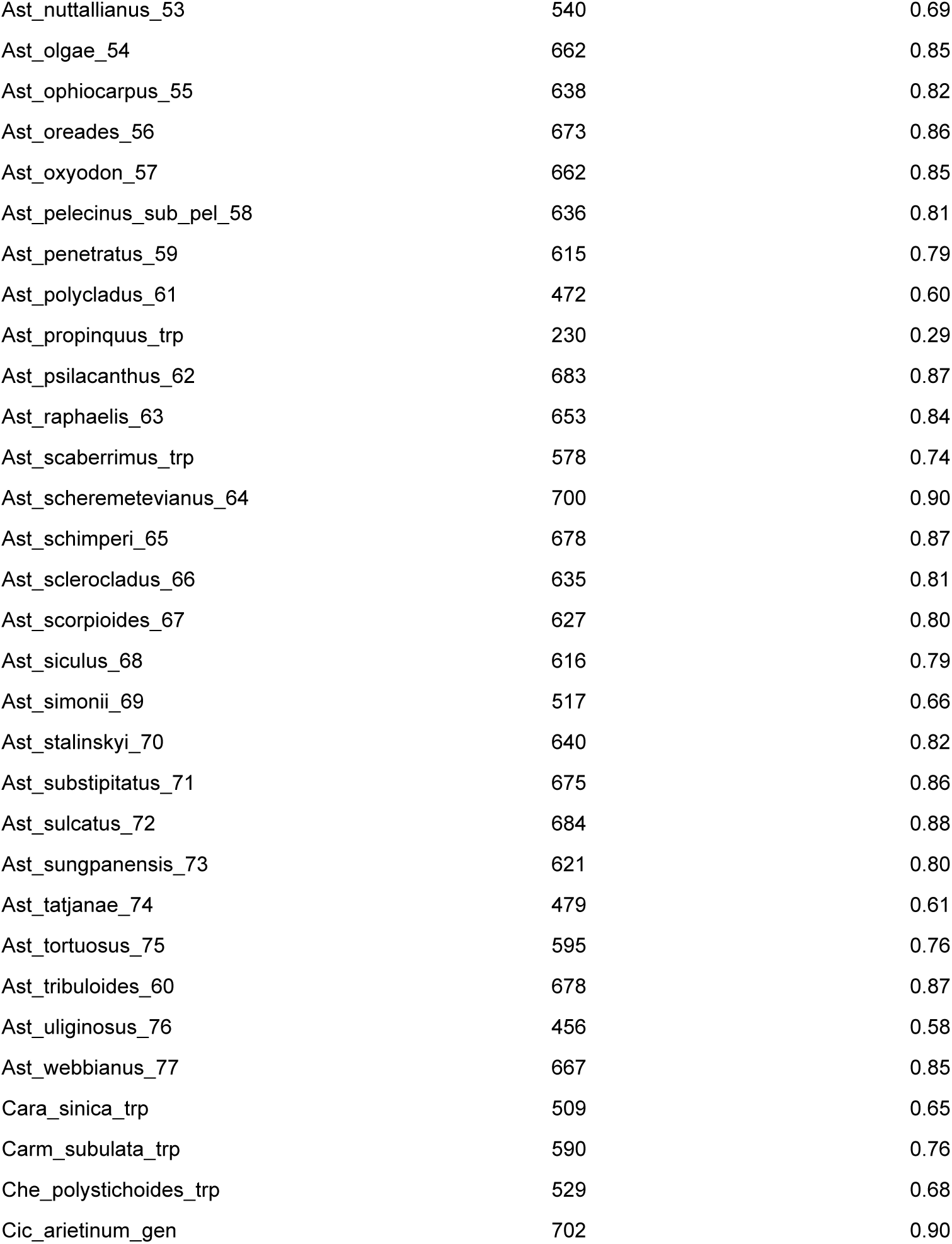
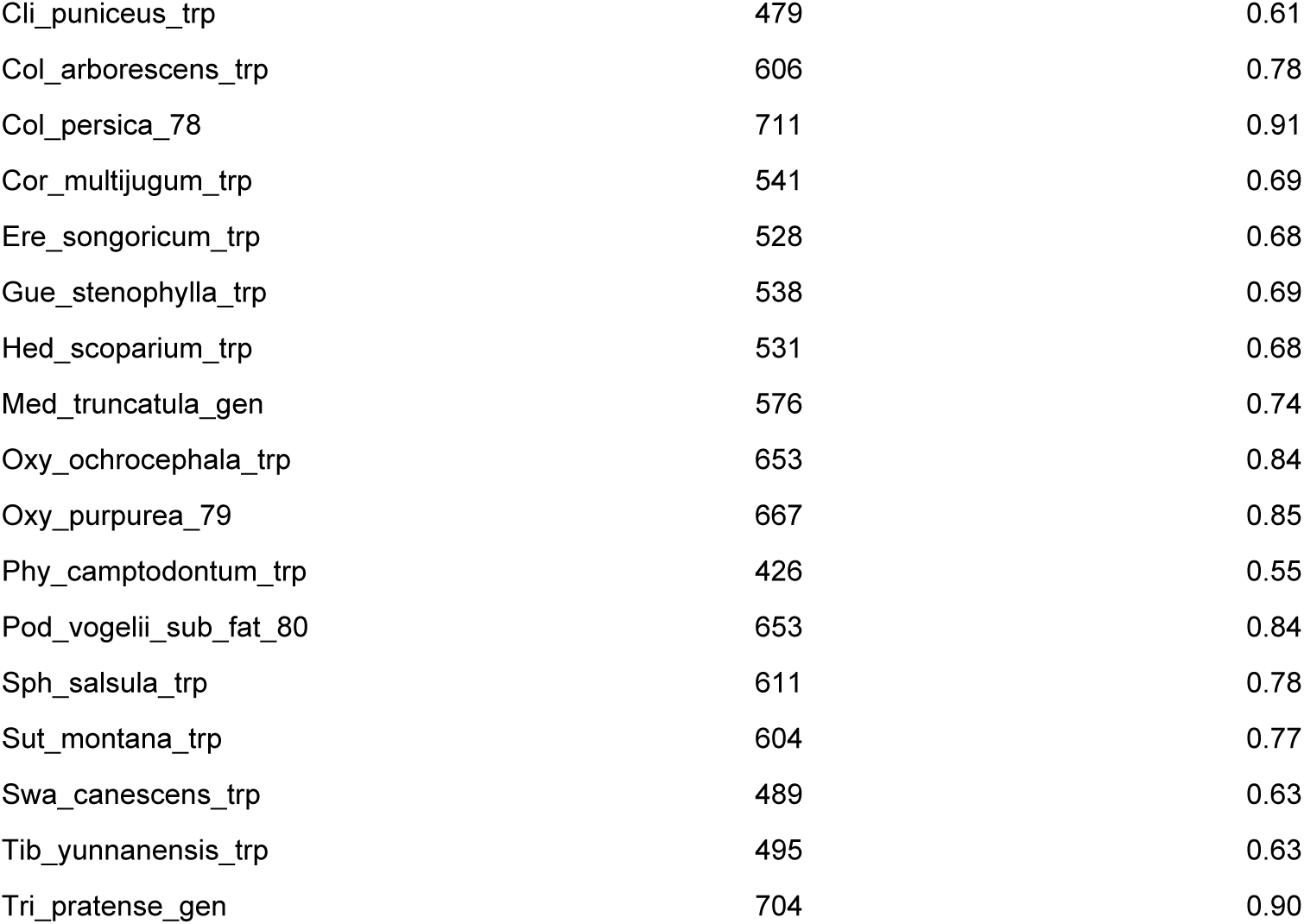
statistics of the orthology inference performed with the monophyletic outgroup (MO) method described by Yang and Smith (2014)

| taxonID | num_ortho | %ortho_out_of_total_781 | AVG_num_ortho_per_spec | range_min-max |
| --- | --- | --- | --- | --- |
| Alh_sparsifolia_trp | 538 | 0.69 |  | 599.8(230-718) |
| Ast_acaulis_1 | 685 | 0.88 |  |  |
| Ast_alopecurus_2 | 669 | 0.86 |  |  |
| Ast_alyssoides_3 | 533 | 0.68 |  |  |
| Ast_annularis_4 | 588 | 0.75 |  |  |
| Ast_apricus_5 | 688 | 0.88 |  |  |
| Ast_arbuscula_trp | 519 | 0.66 |  |  |
| Ast_australis_6 | 643 | 0.82 |  |  |
| Ast_austroaegaeus_7 | 672 | 0.86 |  |  |
| Ast_bhotanensis_trp | 612 | 0.78 |  |  |
| Ast_bisulcatus_var_hay_8 | 539 | 0.69 |  |  |
| Ast_bombycinus_9 | 547 | 0.70 |  |  |
| Ast_brachypetalus_10 | 683 | 0.87 |  |  |
| Ast_canadensis_var_bre_11 | 658 | 0.84 |  |  |
| Ast_caprinus_12 | 698 | 0.89 |  |  |
| Ast_caraganae_13 | 684 | 0.88 |  |  |
| Ast_chlorostachys_14 | 655 | 0.84 |  |  |
| Ast_chorizanthus_15 | 678 | 0.87 |  |  |
| Ast_christianus_sub_chr_16 | 672 | 0.86 |  |  |
| Ast_chrysochlorus_trp | 515 | 0.66 |  |  |
| Ast_contortuplicatus_17 | 681 | 0.87 |  |  |
| Ast_cornu_18 | 506 | 0.65 |  |  |
| Ast_dactylocarpus_19 | 690 | 0.88 |  |  |
| Ast_depressus_20 | 634 | 0.81 |  |  |
| Ast_dipelta_21 | 623 | 0.80 |  |  |
| Ast_dolichopyllus_22 | 610 | 0.78 |  |  |
| Ast_echinatus_23 | 490 | 0.63 |  |  |
| Ast_echinops_24 | 660 | 0.85 |  |  |

|  |  |  |
| --- | --- | --- |
| Ast_epiglottis_25 | 639 | 0.82 |
| Ast_eremophilus_sub_ere_26 | 599 | 0.77 |
| Ast_falconeri_27 | 718 | 0.92 |
| Ast_filicaulis_28 | 580 | 0.74 |
| Ast_frigidus_29 | 704 | 0.90 |
| Ast_fruticosus_30 | 614 | 0.79 |
| Ast_glycyphylloides_31 | 671 | 0.86 |
| Ast_guttatus_32 | 596 | 0.76 |
| Ast_hemiphaca_33 | 543 | 0.70 |
| Ast_hispidulus_34 | 664 | 0.85 |
| Ast_hissaricus_35 | 483 | 0.62 |
| Ast_hoffmeisteri_36 | 707 | 0.91 |
| Ast_hymenostegis_37 | 610 | 0.78 |
| Ast_incanus_38 | 636 | 0.81 |
| Ast_intercedens_39 | 669 | 0.86 |
| Ast_laxmannii_trp | 565 | 0.72 |
| Ast_lentiginosus_trp | 483 | 0.62 |
| Ast_leucocephalus_40 | 627 | 0.80 |
| Ast_lipskyi_41 | 677 | 0.87 |
| Ast_lonchocarpus_42 | 481 | 0.62 |
| Ast_macrobotrys_43 | 655 | 0.84 |
| Ast_macrotropis_44 | 641 | 0.82 |
| Ast_macrourus_45 | 601 | 0.77 |
| Ast_mareoticus_46 | 477 | 0.61 |
| Ast_membranaceus_trp | 286 | 0.37 |
| Ast_microcephalus_47 | 569 | 0.73 |
| Ast_mirabilis_sub_cho_48 | 694 | 0.89 |
| Ast_monspessulanus_49 | 620 | 0.79 |
| Ast_mucronifolius_50 | 582 | 0.75 |
| Ast_norvegicus_51 | 438 | 0.56 |
| Ast_nothoxys_52 | 524 | 0.67 |
| Ast_nuttallianus_53 | 540 | 0.69 |
| Ast_olgae_54 | 662 | 0.85 |
| Ast_ophiocarpus_55 | 638 | 0.82 |
| Ast_oreades_56 | 673 | 0.86 |
| Ast_oxyodon_57 | 662 | 0.85 |
| Ast_pelecinus_sub_pel_58 | 636 | 0.81 |
| Ast_penetratus_59 | 615 | 0.79 |
| Ast_polycladus_61 | 472 | 0.60 |
| Ast_propinquus_trp | 230 | 0.29 |
| Ast_psilacanthus_62 | 683 | 0.87 |
| Ast_raphaelis_63 | 653 | 0.84 |
| Ast_scaberrimus_trp | 578 | 0.74 |
| Ast_scheremetevianus_64 | 700 | 0.90 |
| Ast_schimper_65 | 678 | 0.87 |
| Ast_sclerocladus_66 | 635 | 0.81 |
| Ast_scorpoides_67 | 627 | 0.80 |
| Ast_siculus_68 | 616 | 0.79 |
| Ast_simonii_69 | 517 | 0.66 |
| Ast_stalinskyi_70 | 640 | 0.82 |
| Ast_substipitatus_71 | 675 | 0.86 |
| Ast_sulcatus_72 | 684 | 0.88 |
| Ast_sungpanensis_73 | 621 | 0.80 |
| Ast_tatjanae_74 | 479 | 0.61 |
| Ast_tortuosus_75 | 595 | 0.76 |
| Ast_tribuloides_60 | 678 | 0.87 |
| Ast_uliginosus_76 | 456 | 0.58 |
| Ast_webbianus_77 | 667 | 0.85 |
| Cara_sinica_trp | 509 | 0.65 |
| Carm_subulata_trp | 590 | 0.76 |
| Che_polystichoides_trp | 529 | 0.68 |
| Cic_arietinum_gen | 702 | 0.90 |
| Cli_puniceus_trp | 479 | 0.61 |
| Col_arborescens_trp | 606 | 0.78 |
| Col_persica_78 | 711 | 0.91 |
| Cor_multijugum_trp | 541 | 0.69 |
| Ere_songoricum_trp | 528 | 0.68 |
| Gue_stenophylla_trp | 538 | 0.69 |
| Hed_scoparium_trp | 531 | 0.68 |
| Med_truncatula_gen | 576 | 0.74 |
| Oxy_ochrocephala_trp | 653 | 0.84 |
| Oxy_purpurea_79 | 667 | 0.85 |
| Phy_camptodontum_trp | 426 | 0.55 |
| Pod_vogelii_sub_fat_80 | 653 | 0.84 |
| Sph_salsula_trp | 611 | 0.78 |
| Sut_montana_trp | 604 | 0.77 |
| Swa_canescens_trp | 489 | 0.63 |
| Tib_yunnanensis_trp | 495 | 0.63 |
| Tri_pratense_gen | 704 | 0.90 |

## Notes

### Competing Interest Statement

The authors have declared no competing interest.

## References

Andermann, T., Torres Jiménez, M. F., Matos-Maraví, P., Batista, R., Blanco-Pastor, J. L., Gustafsson, A. L. S., … & Antonelli, A. (2020). A guide to carrying out a phylogenomic target sequence capture project. Frontiers in genetics, 10, 1407.

Andrews, S. (2010). FastQC: A Quality Control Tool for High Throughput Sequence Data [Online]. Available online at: http://www.bioinformatics.babraham.ac.uk/projects/fastqc/

Azani, N., Bruneau, A., Wojciechowski, M. F., & Zarre, S. (2017). Molecular phylogenetics of annual *Astragalus* (Fabaceae) and its systematic implications. Botanical Journal of the Linnean Society, 184(3), 347–365.

Azani, N., Bruneau, A., Wojciechowski, M. F., & Zarre, S. (2019). Miocene climate change as a driving force for multiple origins of annual species in *Astragalus* (Fabaceae, Papilionoideae). Molecular Phylogenetics and Evolution, 137, 210–221.

Barneby, R. C. (1964). Atlas of North American Astragalus: The cercidothrix, hypoglottis, piptoloboid, trimeniaeus, and orophaca Astragali (Vol. 13). New York Botanical Garden.

Bartha, L., Dragoş, N., Molnár V, A., & Sramkó, G. (2013). Molecular evidence for reticulate speciation in *Astragalus* (Fabaceae) as revealed by a case study from sect. Dissitiflori. Botany, 91(10), 702–714.

Bolger, A. M., Lohse, M., & Usadel, B. (2014). Trimmomatic: A flexible trimmer for Illumina Sequence Data. *Bioinformatics*, btu170.

Brown, J. W., Walker, J. F., & Smith, S. A. (2017). Phyx: phylogenetic tools for unix. Bioinformatics, 33(12), 1886–1888.

Chamala, S., García, N., Godden, G. T., Krishnakumar, V., Jordon-Thaden, I. E., De Smet, R., … & Soltis, P. S. (2015). MarkerMiner 1.0: A new application for phylogenetic marker development using angiosperm transcriptomes. Applications in Plant Sciences, 3(4), 1400115.

De Vega, J. J., Ayling, S., Hegarty, M., Kudrna, D., Goicoechea, J. L., Ergon, Å., … & Skøt, L. (2015). Red clover (*Trifolium pratense* L.) draft genome provides a platform for trait improvement. Scientific reports, 5(1), 17394.

Ewels, P., Magnusson, M., Lundin, S., & Käller, M. (2016). MultiQC: summarize analysis results for multiple tools and samples in a single report. Bioinformatics, 32(19), 3047–3048.

Faircloth, B. C., McCormack, J. E., Crawford, N. G., Harvey, M. G., Brumfield, R. T., & Glenn, T. C. (2012). Ultraconserved elements anchor thousands of genetic markers spanning multiple evolutionary timescales. Systematic biology, 61(5), 717–726.

Forrest, L. L., Hart, M. L., Hughes, M., Wilson, H. P., Chung, K. F., Tseng, Y. H., & Kidner, C. A. (2019). The limits of Hyb-Seq for herbarium specimens: impact of preservation techniques. Frontiers in ecology and evolution, 7, 439.

Folk, R. A., Charboneau, J. L., Belitz, M., Singh, T., Kates, H. R., Soltis, D. E., … & Siniscalchi, C. M. (2024). Anatomy of a mega-radiation: Biogeography and niche evolution in *Astragalus*. American Journal of Botany, 111(3), e16299.

González-Domínguez, J., & Schmidt, B. (2016). ParDRe: faster parallel duplicated reads removal tool for sequencing studies. Bioinformatics, 32(10), 1562–1564.

Hardion, L., Dumas, P. J., Abdel-Samad, F., Kharrat, M. B. D., Surina, B., Affre, L., … & Baumel, A. (2016). Geographical isolation caused the diversification of the Mediterranean thorny cushion-like *Astragalus* L. sect. Tragacantha DC.(Fabaceae). Molecular phylogenetics and evolution, 97, 187–195.

Johnson, M. G., Gardner, E. M., Liu, Y., Medina, R., Goffinet, B., Shaw, A. J., … & Wickett, N. J. (2016). HybPiper: Extracting coding sequence and introns for phylogenetics from high-throughput sequencing reads using target enrichment. Applications in plant sciences, 4(7), 1600016.

Katoh, K., & Standley, D. M. (2013). MAFFT multiple sequence alignment software version 7: improvements in performance and usability. Molecular biology and evolution, 30(4), 772–780.

Kazemi, M., Kazempour Osaloo, S., Asghar Maassoumi, A., & Rastegar Pouyani, E. (2009). Molecular phylogeny of selected Old World *Astragalus* (Fabaceae): incongruence among chloroplast trnL-F, ndhF and nuclear ribosomal DNA ITS sequences. Nordic Journal of Botany, 27(5), 425–436.

Kazempour Osaloo, S., Maassoumi, A. A., & Murakami, N. (2005). Molecular systematics of the Old World *Astragalus* (Fabaceae) as inferred from nrDNA ITS sequence data. Brittonia, 57(4), 367–381.

Kazempour Osaloo, S., Maassoumi, A. A., & Murakami, N. (2003). Molecular systematics of the genus *Astragalus* L.(Fabaceae): Phylogenetic analyses of nuclear ribosomal DNA internal transcribed spacers and chloroplast gene ndh F sequences. Plant Systematics and Evolution, 242, 1–32.

Kenicer, G. (2005). Legumes of the World. Edited by G. Lewis, B. Schrire, B. MacKinder & M. Lock. Royal Botanic Gardens, Kew. 2005. xiv+ 577pp., colour photographs & line drawings. ISBN 1 900 34780 6.£ 55.00 (hardback). Edinburgh journal of botany, 62(3), 195–196.

Koenen, E. J. M., De Vos, J. M., Atchison, G. W., Simon, M. F., Schrire, B. D., De Souza, E. R., … & Hughes, C. E. (2013). Exploring the tempo of species diversification in legumes. South African Journal of Botany, 89, 19–30.

Maassoumi, A. A., & Ashouri, P. (2022). The hotspots and conservation gaps of the mega genus *Astragalus* (Fabaceae) in the Old-World. Biodiversity and Conservation, 31(8), 2119–2139.

Maassoumi, A. A., & Khajoei Nasab, F. (2023). Richness and endemism centers of mega genus *Astragalus* (Fabaceae) in Iran. Collectanea Botanica, 42, e001–e001.

Mai, U., & Mirarab, S. (2018). TreeShrink: fast and accurate detection of outlier long branches in collections of phylogenetic trees. BMC genomics, 19, 23–40.

Maylandt, C., Seidl, A., Kirschner, P., Pfanzelt, S., Király, G., Neuffer, B., … & Tremetsberger, K. (2024). Phylogeography of the Euro-Siberian steppe plant *Astragalus austriacus*: Late Pleistocene climate fluctuations fuelled formation and expansion of two main lineages from a Pontic-Pannonian area of origin. Perspectives in Plant Ecology, Evolution and Systematics, 125800.

McKain, M., and M. Wilson. 2017. mrmckain/Fast-Plast: Fast-Plast v.1.2.6 (Version v.1.2.6). Zenodo. 10.5281/zenodo.973887 [accessed 1 February 2024]

McKain, M. R., Johnson, M. G., Uribe-Convers, S., Eaton, D., & Yang, Y. (2018). Practical considerations for plant phylogenomics. Applications in plant sciences, 6(3), e1038.

Minh, B. Q., Schmidt, H. A., Chernomor, O., Schrempf, D., Woodhams, M. D., Von Haeseler, A., & Lanfear, R. (2020). IQ-TREE 2: new models and efficient methods for phylogenetic inference in the genomic era. Molecular biology and evolution, 37(5), 1530–1534.

Moonlight, P. W., Baldaszti, L., Cardoso, D., Elliott, A., Särkinen, T., & Knapp, S. (2024). Twenty years of big plant genera. Proceedings of the Royal Society B, 291(2023), 20240702.

Morales-Briones, D. F., Kadereit, G., Tefarikis, D. T., Moore, M. J., Smith, S. A., Brockington, S. F., … & Yang, Y. (2021). Disentangling sources of gene tree discordance in phylogenomic data sets: testing ancient hybridizations in Amaranthaceae sl. Systematic Biology, 70(2), 219–235.

Morales-Briones, D.F., B. Gehrke, H. Chien-Hsun Huang, A. Liston, M. Hong. H.E. Marx, D.C. Tank & Y. Yang. 2022. Analysis of paralogs in target enrichment data pinpoints multiple ancient polyploidy events in *Alchemilla* s.l. (Rosaceae). Systematic Biology 71(1):190–207

Pease, J. B., Brown, J. W., Walker, J. F., Hinchliff, C. E., & Smith, S. A. (2018). Quartet sampling distinguishes lack of support from conflicting support in the green plant tree of life. American journal of botany, 105(3), 385–403.

Pezzini, F. F., Ferrari, G., Forrest, L. L., Hart, M. L., Nishii, K., & Kidner, C. A. (2023). Target capture and genome skimming for plant diversity studies. Applications in Plant Sciences, 11(4), e11537.

Podlech D., Zarre S. (2013). A taxonomic revision of the genus Astragalus L. (Leguminosae) in the Old World. Vienna: Naturhistorisches Museum.

POWO (2024). Plants of the World Online. Facilitated by the Royal Botanic Gardens, Kew. Published on the Internet; https://powo.science.kew.org/. Retrieved 04 September 2024.

Prjibelski, A., Antipov, D., Meleshko, D., Lapidus, A., & Korobeynikov, A. (2020). Using SPAdes de novo assembler. Current protocols in bioinformatics, 70(1), e102.

Ranwez, V., Douzery, E. J., Cambon, C., Chantret, N., & Delsuc, F. (2018). MACSE v2: toolkit for the alignment of coding sequences accounting for frameshifts and stop codons. Molecular biology and evolution, 35(10), 2582–2584.

Rundel, P. W., Huggins, T. R., Prigge, B. A., & Rasoul Sharifi, M. (2015). Rarity in *Astragalus*: a California perspective. Aliso: A Journal of Systematic and Floristic Botany, 33(2), 111–120.

Sanderson, M. J., & Wojciechowski, M. F. (1996). Diversification rates in a temperate legume clade: are there “so many species” of *Astragalus* (Fabaceae)?. American Journal of Botany, 83(11), 1488–1502.

Scherson, R. A., Vidal, R., & Sanderson, M. J. (2008). Phylogeny, biogeography, and rates of diversification of New World *Astragalus* (Leguminosae) with an emphasis on South American radiations. American Journal of Botany, 95(8), 1030–1039.

Smith, S. A., Moore, M. J., Brown, J. W., & Yang, Y. (2015). Analysis of phylogenomic datasets reveals conflict, concordance, and gene duplications with examples from animals and plants. BMC evolutionary biology, 15, 1–15.

Soltani, E., Benakashani, F., Baskin, J. M., & Baskin, C. C. (2021). Reproductive biology, ecological life history/demography and genetic diversity of the megagenus *Astragalus* (Fabaceae, Papilionoideae). The Botanical Review, 87, 55–106.

Soltis, D. E., & Kuzoff, R. K. (1995). Discordance between nuclear and chloroplast phylogenies in the Heuchera group (Saxifragaceae). Evolution, 49(4), 727–742.

Stamatakis, A. (2014). RAxML Version 8: A tool for Phylogenetic Analysis and Post-Analysis of Large Phylogenies. Bioinformatics, open access link: http://bioinformatics.oxfordjournals.org/content/early/2014/01/21/bioinformatics.btu033.abstract?keytype=ref&ijkey=VTEqgUJYCDcf0kP

Su, C., Duan, L., Liu, P., Liu, J., Chang, Z., & Wen, J. (2021). Chloroplast phylogenomics and character evolution of eastern Asian *Astragalus* (Leguminosae): Tackling the phylogenetic structure of the largest genus of flowering plants in Asia. Molecular Phylogenetics and Evolution, 156, 107025.

Tang, H., Krishnakumar, V., Bidwell, S., Rosen, B., Chan, A., Zhou, S., … & Town, C. D. (2014). An improved genome release (version Mt4. 0) for the model legume *Medicago truncatula*. BMC genomics, 15, 1–14.

Vargas, O. M., Heuertz, M., Smith, S. A., & Dick, C. W. (2019). Target sequence capture in the Brazil nut family (Lecythidaceae): Marker selection and in silico capture from genome skimming data. Molecular Phylogenetics and Evolution, 135, 98–104.

Varshney, R. K., Song, C., Saxena, R. K., Azam, S., Yu, S., Sharpe, A. G., … & Cook, D. R. (2013). Draft genome sequence of chickpea (*Cicer arietinum*) provides a resource for trait improvement. Nature biotechnology, 31(3), 240–246.

Vatanparast, M., Powell, A., Doyle, J. J., & Egan, A. N. (2018). Targeting legume loci: A comparison of three methods for target enrichment bait design in Leguminosae phylogenomics. Applications in Plant Sciences, 6(3), e1036.

Wen, D., Yu, Y., Zhu, J., & Nakhleh, L. (2018). Inferring phylogenetic networks using PhyloNet. Systematic biology, 67(4), 735–740.

Wojciechowski, M. F. (2005). *Astragalus* (Fabaceae): A molecular phylogenetic perspective. Brittonia, 57(4), 382–396.

Wojciechowski, M. F., Sanderson, M. J., & Hu, J. M. (1999). Evidence on the monophyly of *Astragalus* (Fabaceae) and its major subgroups based on nuclear ribosomal DNA ITS and chloroplast DNA trnL intron data. Systematic Botany, 409–437.

Wojciechowski, M. F., Sanderson, M. J., Baldwin, B. G., & Donoghue, M. J. (1993). Monophyly of aneuploid *Astragalus* (Fabaceae): evidence from nuclear ribosomal DNA internal transcribed spacer sequences. American Journal of Botany, 80(6), 711–722.

Yang, Y., & Smith, S. A. (2014). Orthology inference in nonmodel organisms using transcriptomes and low-coverage genomes: improving accuracy and matrix occupancy for phylogenomics. Molecular biology and evolution, 31(11), 3081–3092.

Yu, Y., & Nakhleh, L. (2015). A maximum pseudo-likelihood approach for phylogenetic networks. BMC genomics, 16, 1–10.

Zarre, S., & Azani, N. (2013). Perspectives in taxonomy and phylogeny of the genus *Astragalus* (Fabaceae): a review. Progress in biological sciences, 3(1), 1–6.

Záveská, E., Maylandt, C., Paun, O., Bertel, C., Frajman, B., Schönswetter, P., & Steppe Consortium. (2019). Multiple auto-and allopolyploidisations marked the Pleistocene history of the widespread Eurasian steppe plant *Astragalus* onobrychis (Fabaceae). Molecular Phylogenetics and Evolution, 139, 106572.

Zhang, C., Scornavacca, C., Molloy, E. K., & Mirarab, S. (2020). ASTRAL-Pro: quartet-based species-tree inference despite paralogy. Molecular biology and evolution, 37(11), 3292–3307.

Zhao, Y., Zhang, R., Jiang, K. W., Qi, J., Hu, Y., Guo, J., Zhu, R., Zhang, T., Egan, A. N., Yi, T., Huang, C., & Ma, H. (2021). Nuclear phylotranscriptomics and phylogenomics support numerous polyploidization events and hypotheses for the evolution of rhizobial nitrogen-fixing symbiosis in Fabaceae. Molecular Plant, 14(5), 748–773.

Zimmer, E. A., & Wen, J. (2015). Using nuclear gene data for plant phylogenetics: Progress and prospects II. Next-gen approaches. Journal of Systematics and Evolution, 53(5), 371–379.

